# A Multi-Disciplinary Framework for Decoding S1PR1-Selective Agonism

**DOI:** 10.1101/2025.08.27.672536

**Authors:** Leiye Yu, Haizhan Jiao, Bin Pang, Rujuan Ti, Bing Gan, Zhaoyang Qin, Jinxin Wang, Lizhe Zhu, Hongli Hu, Ruobing Ren

**Affiliations:** Shanghai Key Laboratory of Metabolic Remodeling and Health, Institute of Metabolism and Integrative Biology, Shanghai Xuhui Central Hospital, Zhongshan-Xuhui Hospital, Fudan University, Shanghai 200437, China; Department of Otorhinolaryngology Head and Neck Surgery, Shanghai Sixth People’s Hospital Affiliated to Shanghai Jiao Tong University School of Medicine, Shanghai 200233, China; Kobilka Institute of Innovative Drug Discovery, School of Life and Health Sciences, The Chinese University of Hong Kong, Shenzhen 518172, China; Warshal Institute of Computational Biology, School of Life and Health Sciences, The Chinese University of Hong Kong, Shenzhen 518172, China; School of Pharmacy, Second Military Medical University, Shanghai, China

**Keywords:** GPCR, S1P, S1PR1, CryoEM, Selective agonism

## Abstract

Sphingosine-1-phosphate (S1P), a key metabolite of sphingolipids, plays crucial roles in a wide range of physiological and pathological processes. S1P primarily exerts its functions by binding to G protein-coupled sphingosine-1-phosphate receptors (S1PRs), which include five subtypes (S1PR1–5), thereby activating these receptors and their downstream signaling pathways. Understanding the molecular determinants that govern agonist selectivity among different S1PR subtypes is vital for the rational and precise development of targeted therapeutic agents. Here, four cryo-electron microscopy structures of agonist-bound S1PR1-Gi complexes are reported. Through an integrated approach combining structural analysis, molecular dynamics simulations, and pharmacological assays, the molecular basis for the selectivity of CYM5442, HY-X-1011, Ponesimod, and SAR247799 toward S1PR1 over S1PR3 and S1PR5 is uncovered. Specifically, the selectivity arises from a combination of non-conserved residues within the ligand-binding pocket and at the Gi-protein interface, distinct curved agonist-binding modes oriented toward transmembrane helices 5-7 that cause steric clashes with S1PR3, and the presence of branched moieties at the lower part of three agonists. These features collectively enhance agonist potency and efficacy for S1PR1 while reducing activity at S1PR3 and S1PR5. These findings establish a structural framework for the rational design of next-generation S1PR1 highly selective agonists with improved therapeutic potential.

## 1. Introduction

Sphingosine-1-phosphate (S1P), a key metabolite of sphingolipids, plays crucial roles in a wide range of physiological processes, including lymphocyte trafficking and cardiovascular development. S1P is generated through the phosphorylation of sphingosine, a backbone structural component of all sphingolipids, and can be exported from cells via specific membrane transporters^1–4^. S1P primarily exerts its biological functions by binding to G protein-coupled sphingosine-1-phosphate receptors (S1PRs), which consist of five subtypes (S1PR1–5), thereby activating these receptors and triggering various downstream signaling pathways.^5^ S1PRs belong to the class A GPCR (G protein-coupled receptor) superfamily. Activation of GPCRs involves significant conformational rearrangements on the cytoplasmic side, particularly a large outward movement of transmembrane helix 6 (TM6), accompanied by reorganization of other helices, which creates an intracellular pocket that can engage different G proteins (Gi/o, Gs, Gq, and G12/13), GRKs, and arrestins to form functional signaling complexes^6^. The S1PR subtypes activate distinct yet partially overlapping G protein-mediated signaling cascades, thus orchestrating diverse cellular responses such as proliferation, apoptosis, adhesion, and migration. Specifically, S1PR1 couples exclusively with Gi/o (the alpha subunit of heterotrimeric G proteins), S1PR2–3 interact with Gi/o, G12/13, and Gq, while S1PR4–5 bind to Gi/o and G12/13^5^.

S1PR1 is highly expressed in endothelial cells and adipocytes, with lower expression levels observed in various immune cell populations. Upon activation by S1P, S1PR1 engages Gi proteins, leading to subsequent activation of Rac1 signaling, which promotes the formation of tight junctions between endothelial cells and helps maintain the integrity and normal function of vascular and lymphatic endothelial barriers^7^. In vascular endothelial cells (VECs), S1PR1 signaling not only stabilizes adherens junctions but also enhances nitric oxide (NO) production via endothelial nitric oxide synthase (eNOS), which is essential for regulating blood flow and pressure^8^. Moreover, S1PR1 plays a pivotal role in lymphocyte egress from the thymus and secondary lymphoid organs, a process that relies on a concentration gradient of S1P-high in plasma (∼1 μM) and lymph (∼100 nM), compared to much lower levels in interstitial fluids^9–13^. Additionally, S1PR1 is expressed in brain endothelial cells, astrocytes, glial cells, and leukocytes, where it contributes significantly to the proper functioning of the central nervous system.

S1PRs play complex roles in multiple pathological processes, including cardiovascular, autoimmune, inflammatory, neurological, oncologic, hearing loss, and fibrotic diseases ^14^. Consequently, the therapeutic potential of S1PR1 modulation has been extensively investigated, leading to the development of numerous S1PR1 modulators, several of which are clinically approved pharmacological agents for treating various conditions such as multiple sclerosis (MS) and inflammatory bowel disease (IBD)^15–18^. Multiple sclerosis (MS) is a chronic neurodegenerative disorder characterized by persistent inflammation within the central nervous system (CNS), resulting in neuronal demyelination and subsequent functional impairments. Fingolimod (FTY720) was the first FDA-approved drug specifically targeting S1PR1 for the treatment of MS^16^. Fingolimod undergoes phosphorylation in vivo to form phosphorylated Fingolimod (FTY720-P). Upon binding to S1PR1, FTY720-P functionally antagonizes the receptor via β-arrestin-mediated internalization, significantly reducing circulating and CNS-infiltrating immune cell populations, thereby suppressing inflammation and disease progression. However, patients treated with Fingolimod may experience adverse events such as relapse of multiple sclerosis, decreased lymphocyte count, fatigue, and headache^19^. Furthermore, FTY720-P also activates all five S1PR subtypes and engages G protein-dependent signaling pathways that can promote immune cell migration^20^. Additionally, its long half-life contributes to further side effects^21^. In recent years, the FDA has approved three second-generation S1PR-targeted drugs—Siponimod, Ozanimod, and Ponesimod—for the treatment of MS ^22–24^. Although these newer agents exhibit improved receptor selectivity, they still carry risks of side effects such as bradycardia. Siponimod and Ozanimod selectively activate both S1PR1 and S1PR5 with similar potency, with Siponimod showing approximately a thousand-fold lower potency toward S1PR3 and S1PR4 compared to S1PR1 and S1PR5^25,26^. Ponesimod strongly activates S1PR1 but exhibits lower affinity for S1PR3 and S1PR5^27^. The activation of multiple S1PR subtypes may lead to varied physiological effects and increase the likelihood of certain adverse reactions, such as bradycardia^28^.

It is noteworthy that current therapeutics predominantly target S1PR1 as their primary mechanism of action. However, due to the high sequence and structural similarity among the five S1PR subtypes, existing modulators often display cross-reactivity across multiple receptor subtypes, activating diverse signaling pathways. This phenomenon not only complicates therapeutic outcomes but also contributes to the potential side effects associated with these drugs. Therefore, developing subtype-selective S1PR modulators remains a significant challenge. Understanding the molecular determinants that govern agonist selectivity among S1PR subtypes is crucial for the rational and precise design of targeted therapies. While multiple structures of S1PRs bound to various modulators have been reported previously^29–37^, the mechanisms underlying agonist selectivity among different S1PR subtypes remain insufficiently understood. In this study, we determined four agonist-bound S1PR1-Gi complex structures. Through integrated structural analysis, mutagenesis combined with Gi coupling assays, and molecular dynamics simulations, we uncovered the molecular basis for the preferential selectivity of agonists toward S1PR1 over S1PR3 and S1PR5. These findings provide a foundation for the rational design and development of highly selective S1PR1 modulators.

## 2. Results

### 2.1 Overall structures of the S1PR1-Gi complexes bound with CYM5442, HY-X-10111, Ponesimod, or SAR247799

Previously, several agonists—CYM5442, HY-X-1011, Ponesimod, and SAR247799—were reported to exhibit improved selectivity for S1PR1 over S1PR2–S1PR5 (Figure S1A–D, Supporting Information)^38–41^. Although the activation of S1PR by these four agonists has been characterized previously using various experimental approaches, we employed a standardized bioluminescence resonance energy transfer (BRET)-based assay to systematically evaluate their activation profiles across S1PR1–S1PR5. This method enables quantitative assessment of Gi protein dissociation kinetics and facilitates direct comparison of their pharmacological properties. The BRET assay results revealed that all four agonists exhibit greater potency and efficacy for S1PR1 compared to S1PR3 and S1PR5, with EC50 values differing by one to two orders of magnitude (Fig. 1A–D). Additionally, no activation of S1PR2 or S1PR4 was observed in response to any of the four agonists in the BRET-based Gi dissociation assay (data not shown).

**Figure 1.**
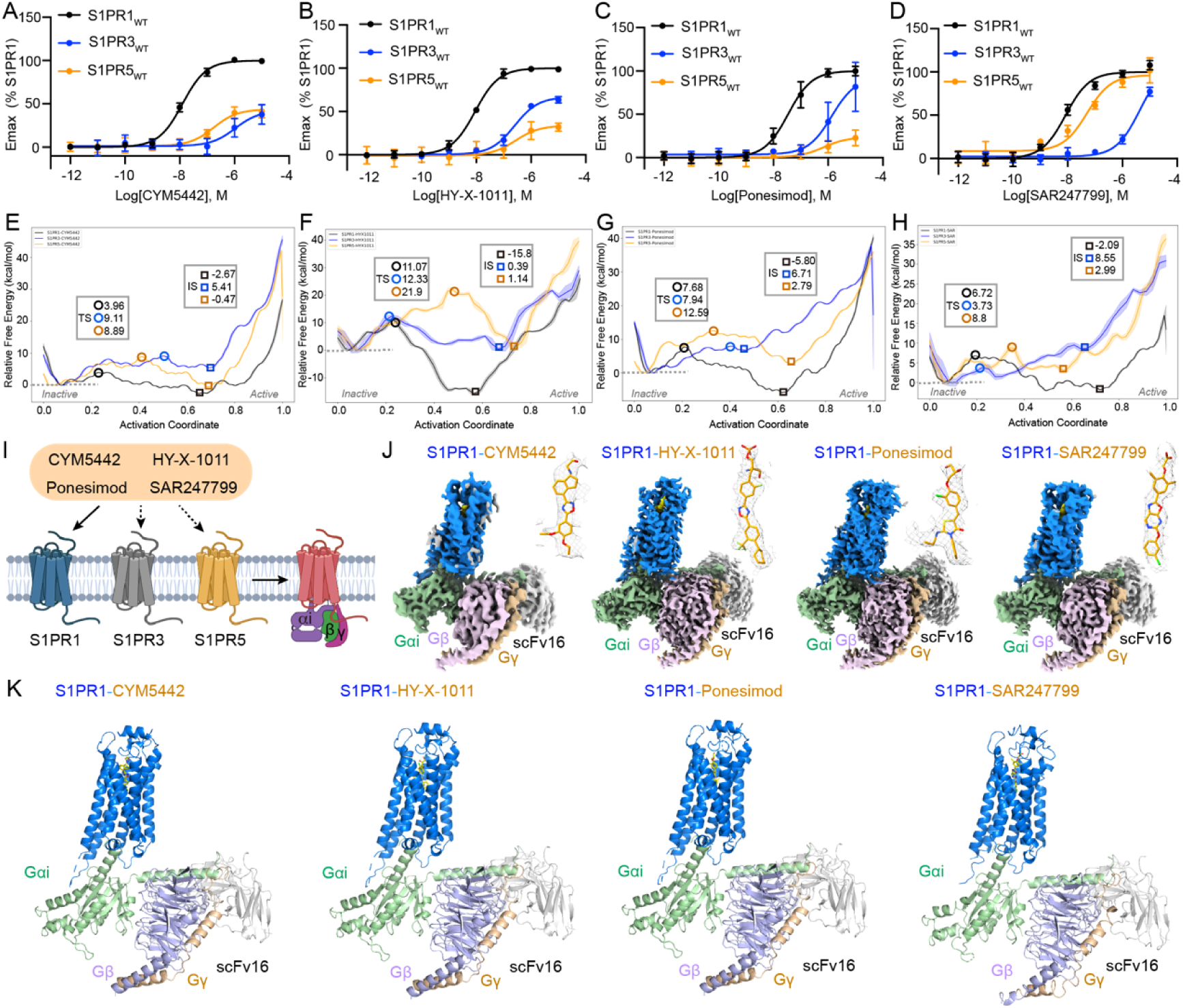
Structures of four S1PR1-Gi complexes bound with CYM5442, HY-X-1011, Ponesimod, or SAR247799. **A-D.** Activation of S1PR1, S1PR3, and S1PR5 by CYM5442, HY-X-1011, Ponesimod, or SAR247799, as measured via the BRET-based Gi dissociation assay. Data are presented as mean ± SEM; n = 3. All four compounds exhibit higher selectivity for S1PR1 compared to S1PR3 and S1PR5. No Gi signaling response is observed for S1PR2 and S1PR4 upon treatment with these agonists (data not shown). **E-H.** Free energy profiles along the activation pathways of S1PR1 (black), S1PR3 (blue), and S1PR5 (wheat) induced by CYM5442, HY-X-1011, Ponesimod, or SAR247799. The profiles are plotted as lines. TS: transition state; IS: intermediate state, structurally resembling the fully active conformation. The free energy differences between the IS/TS states and the inactive state are calculated and labeled using circles (TS) and squares (IS), respectively. **I.** Schematic illustration showing the activation of S1PR1, S1PR3, and S1PR5 by the four agonists—CYM5442, HY-X-1011, Ponesimod, and SAR247799. Solid arrows indicate strong activation, while dashed arrows represent weak activation. **J.** Cryo-EM density maps of S1PR1-Gi complexes bound with CYM5442, HY-X-1011, Ponesimod, or SAR247799. Ligands (yellow) are displayed within the density maps (gray mesh) on the right side of the models and are represented in stick form. **K.** Structural cartoon models of human S1PR1 in complex with Gi, scFv16, and each of the four agonists: CYM5442, HY-X-1011, Ponesimod, or SAR247799. In panels **J–K,** S1PR1 is colored marine, Gαi pale green, Gβ light blue, Gγ wheat, scFv16 gray, and agonists yellow.

To further explore the activation mechanisms, we conducted molecular dynamics simulations using the previously established traveling-salesman-based automated path searching (TAPS) method to identify the minimum free energy path (MFEP) during the transition of S1PR1/3/5 from an inactive to an active conformation upon agonist binding^42^. The activation potential of each ligand can be quantified by the sum of the barrier height ΔG_T_ (the Gibbs free energy difference between the inactive state and the transition state) and the stability of the intermediate state relative to the inactive state ΔG_I_ (the Gibbs free energy difference between the inactive state and the intermediate state; the intermediate state closely resembles the fully active structure)^41^. The calculated free energy distributions (ΔG_T_ + ΔG_I_) along the MFEP for the four distinct ligands and three S1PR subtypes showed strong agreement with the BRET measurements of Gi protein dissociation (Figure 1E–H).

However, the lack of comprehensive structural, computational, and pharmacological profiling has limited our understanding of the mechanisms underlying receptor subtype selectivity (Figure 1I). To investigate and clarify the basis of this selectivity, we determined four Gi-coupled S1PR1 structures bound to CYM5442, HY-X-1011, Ponesimod, or SAR247799 using single-particle cryo-electron microscopy (cryo-EM). The four complexes were successfully assembled following previously established protocols (Figure S1E–H, Supporting Information)9. Data collection and analysis yielded four high-resolution density maps of the S1PR1-Gi complex bound to each respective agonist at overall resolutions of 3.69 Å, 2.79 Å, 2.79 Å, and 2.97 Å, respectively (Figure 2A–E; Table S1, Supporting Information). These maps allowed unambiguous modeling of all seven transmembrane helices of S1PR1 as well as the G protein components (Figure 1J–K; Figure S2F–I, Supporting Information). Importantly, clear electron densities corresponding to each of the four agonists were observed in their respective maps, enabling accurate ligand fitting and detailed characterization of the binding sites (Figure 1J).

**Figure 2.**
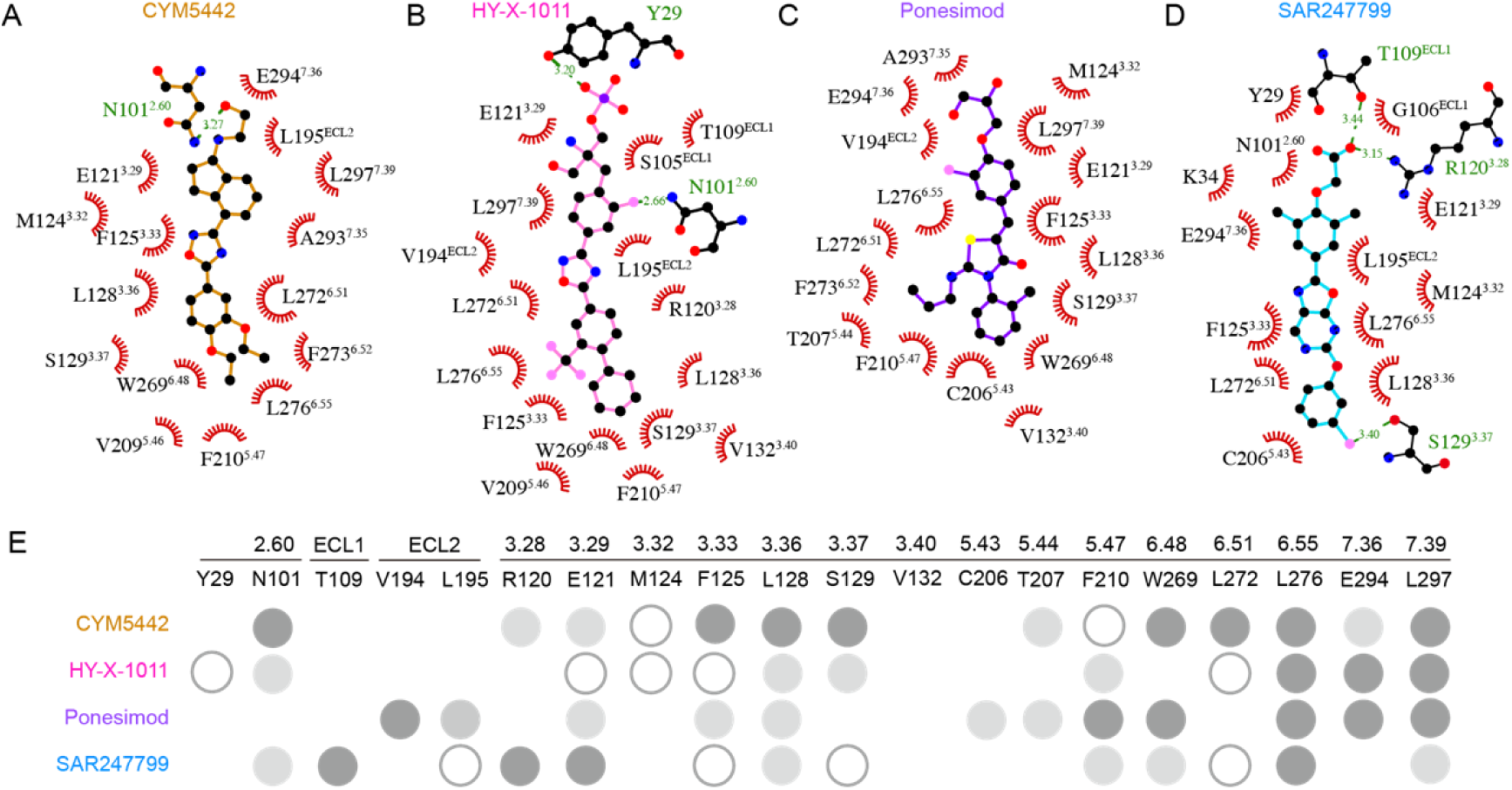
Critical residues involved in the interaction with CYM5442, HY-X-1011, Ponesimod, and SAR247799 that are essential for S1PR1 activation by these four agonists. **A-D.** Residues within the ligand-binding pocket that interact with CYM5442 **(A)**, HY-X-1011 **(B)**, Ponesimod **(C**), or SAR247799 **(D)**. Hydrogen bonds are indicated by green lines, with distances labeled accordingly. **E.** Key binding pocket residues critical for S1PR1 activation mediated by CYM5442, HY-X-1011, Ponesimod, or SAR247799, as confirmed by the BRET-based Gi dissociation assay. In this assay, most binding residues were evaluated through alanine substitution. The effects of residue mutations on the Gi signaling response are represented as follows: white circles with gray borders > dark gray circles > light gray circles. White circles with gray borders indicate that alanine substitution at that position almost completely abolished S1PR1 activation. Corresponding data are provided in Table S2 of the Supporting Information.

### 2.2 Agonist recognition and activation mechanisms of S1PR1

In these Gi-coupled S1PR1 structures, four agonists adopt extended conformations within the orthosteric binding pocket of S1PR1, exhibiting length-dependent variations in polar interactions in the upper portion of the pocket, while maintaining similar hydrophobic contacts in the middle and lower regions (Figure 2A–D). Systematic alanine scanning mutagenesis confirmed the significant roles of most selected residues involved in agonist interaction, as revealed by Gi dissociation BRET assays (Table S2, Supporting Information). These key residues are primarily located in transmembrane segments TM3, TM5, TM6, and TM7 (Figure 2E). Notably, most residues in the middle and bottom regions of the pocket strongly influence the potency and efficacy of all four agonists, including F125³·³³, L128³·³⁶, F210⁵·⁴⁷, L276⁶·⁵⁵, and L297⁷·³⁹. These results suggest that, in addition to the head groups, the hydrophobic tails of these agonists—which interact with TM3, TM5, TM6, and TM7—play comparably critical roles in activating S1PR1 and initiating downstream Gi signaling.

A comparison of conformational changes between agonist-bound active S1PR1 structures and the antagonist ML-056-bound inactive structure provides key mechanistic insights into receptor activation. All active structures display a characteristic outward movement of TM6 by approximately 11.5 Å, which facilitates G protein coupling (Figure S3A–D, Supporting Information). Agonist binding induces a conserved reorientation of side chains in the hydrophobic residues L128³·³⁶, F210⁵·⁴⁷, F273⁵·⁵², W269⁶·⁴⁸, and L297⁷·³⁹ (Figure S3E–H, Supporting Information), with F210⁵·⁴⁷ undergoing a rotamer switch that drives TM6 displacement, accompanied by rotation of F273⁵·⁵². Furthermore, class A GPCR activation-related motifs in S1PR1—including E^3.49^R^3.50^Y^3.51^, P^5.50^I^3.40^F^6.44^, and N^7.49^P^7.50^XXY^7.53^—are largely conserved^43^. These motifs (E141^3.49^R142^3.50^Y143^3.51^, L213^5.50^V132^3.40^F265^6.44^, and N307^7.49^P308^7.50^XXY311^7.53^) exhibit significant conformational changes across all four activated receptor structures (Figure S3I–T, Supporting Information), underscoring the evolutionary conservation of the S1PR activation mechanism.

### 2.3 Impact of non-conserved residues in the receptor pocket on agonist selectivity for S1PR1, S1PR3, and S1PR5

Although most residues that interact with the four agonists are conserved across S1PR1, S1PR3, and S1PR5, we hypothesize that nonconserved residues within the ligand-binding pocket contribute to receptor subtype selectivity. Initially, we identified nonconserved residues in the ligand-binding pockets of S1PR1 and S1PR3 (Figure 3A–D), which are located in the middle and lower regions of the binding cavity. Two previously reported key residues—L276⁶·⁵⁵ and L297⁷·³⁹ in S1PR1, corresponding to F263⁶·⁵⁵ and I284⁷·³⁹ in S1PR3—are known to influence the selectivity of Siponimod and CBP-307 for S1PR1 and S1PR3, respectively. Functional characterization of reciprocal mutants confirmed the importance of these two residue pairs. The L276⁶·⁵⁵F-L297⁷·³⁹I double mutant of S1PR1 exhibited reduced agonist potency and efficacy, similar to or weaker than that of wild-type S1PR3. Conversely, introducing the F263⁶·⁵⁵L-I284⁷·³⁹L mutations into S1PR3 enhanced the potency of CYM5442, HY-X-1011, and Ponesimod to levels approaching those observed for S1PR1, although SAR247799 remained tenfold less potent (Figure 3E–H, Table S3, Supporting Information). Additionally, residues S129³·³⁷, T207⁵·⁴⁴, and V132³·⁴⁰ in S1PR1 correspond to G123³·³⁷, I201⁵·⁴⁴, and T126³·⁴⁰ in S1PR3, and all contact the lower portions of the four agonists. Therefore, we examined the effects of mutating these three residues pairs—S129³·³⁷G/G123³·³⁷S, T207⁵·⁴⁴I/I201⁵·⁴⁴T, and V132³·⁴⁰T/T126³·⁴⁰V—on receptor activation. These mutants displayed partial exchange effects in response to Ponesimod and SAR247799, except for V132³·⁴⁰T in S1PR1 and T126³·⁴⁰V in S1PR3, which had no observable impact (Figure S4A–F, Supporting Information). Three non-conserved residues—F133³·⁴¹, V209⁵·⁴⁶, and L213⁵·⁵⁰—located at the bottom of the S1PR1 pocket show distinct interaction patterns. Among them, V209⁵·⁴⁶ interact with the tail ends of CYM5442, HY-X-1011, and Ponesimod. In contrast, F133³·⁴¹, and L213⁵·⁵⁰ do not exhibit direct interactions with any of the four agonists in our structural analyses. Intriguingly, results from the Gi dissociation assay showed that triple mutants F133³·⁴¹C-V209⁵·⁴⁶I-L213⁵·⁵⁰I (S1PR1) and C127³·⁴¹F-I203⁵·⁴⁶V-I207⁵·⁵⁰L (S1PR3) exhibited exchanged activation potencies when stimulated by CYM5442, HY-X-1011, Ponesimod, and SAR247799 (Figure 3I–L, Table S3, Supporting Information).

**Figure 3.**
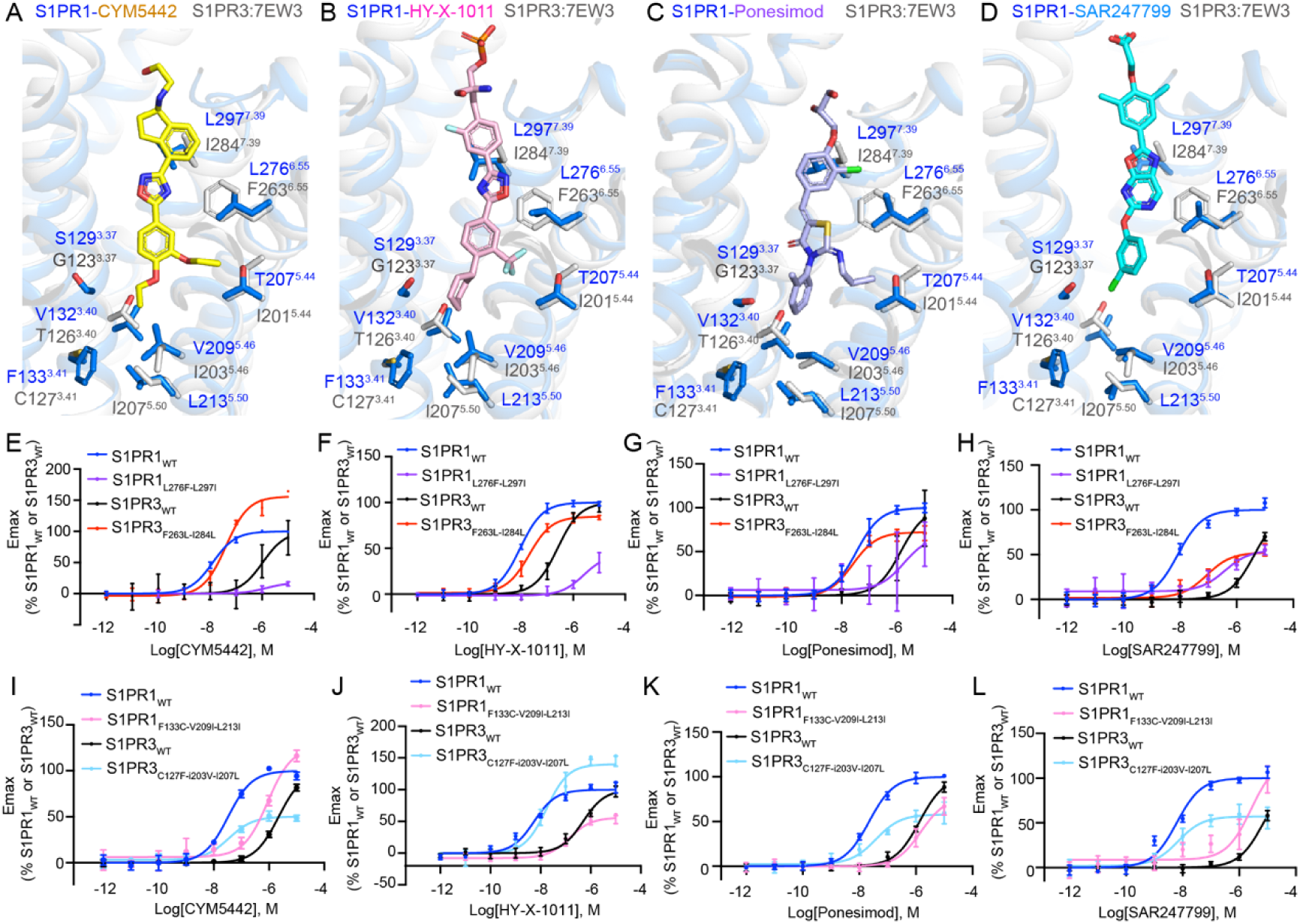
Nonconserved residues influence the selectivity of four agonists for S1PR1 and S1PR3. **A-D.** Nonconserved residues in the ligand binding pocket of S1PR1 and S1PR3. CYM5442-, HY-X-1011-, Ponesimod-, or SAR247799-bound S1PR1 are superimposed with S1PR3 (PDB: 7EW3) in panels **A-D,** respectively. Residues of S1PR1 (marine) and S1PR3 (gray) are shown as sticks. Molecules are shown as sticks and colored yellow (CYM5442), pink (HY-X-1011), light blue (Ponesimod), and cyan (SAR247799). **E-L.** BRET-based Gi dissociation assays showing activation of swapped mutants of S1PR1 and S1PR3 by the four agonists. Data are presented as mean ± SEM; n=3.

Similarly, we also identified several residues predominantly clustered in the central and lower regions of the binding pockets of S1PR1 and S1PR5 (Figure 4A–D). Among them, M124^3.32^ and S129^3.37^ (S1PR1), corresponding to V115^3.32^ and T120^3.37^ (S1PR3), exhibit close contacts with four agonists. However, these reciprocal mutants exhibit varying effects on the activation of S1PR1 and S1PR5 by the four agonists (Figure S5A–H, Supporting Information). We further analyzed T207^5.44^ and V198^5.44^, which are located between TM5 and TM6, near the branched moieties at the lower portions of CYM5442, HY-X-1011, and Ponesimod. The T207^5.44^V mutation in S1PR1 results in reduced potency, similar to the activation of S1PR3 by the four agonists, whereas the V198^5.44^T mutation in S1PR3 leads to increased potency, resembling the activation of S1PR1 by CYM5442, HY-X-1011, and SAR247799 (Figure 4E–H). Additionally, three residues—F133³·⁴¹, V209⁵·⁴⁶, and L213⁵·⁵⁰—at the bottom of the S1PR1 pocket are also not conserved in S1PR5 (L127³·⁴¹, A203⁵·⁴⁶, I207⁵·⁵⁰) (Figure 4A–D). Reciprocal triple mutations—F133³·⁴¹L-V209⁵·⁴⁶A-L213⁵·⁵⁰I in S1PR1 and L127³·⁴¹F-A203⁵·⁴⁶V-I207⁵·⁵⁰L in S1PR5—resulted in fully swapped activation profiles for CYM5442 and SAR247799, and showed significant exchange in activation effects for Ponesimod and HY-X-1011, except for L127³·⁴¹F-A203⁵·⁴⁶V-I207⁵·⁵⁰L in S1PR5 in response to Ponesimod (Figure 4I–L). Collectively, our findings systematically reveal nonconserved residue pairs within the ligand-binding pocket that contribute to subtype selectivity of S1PR1 by the four agonists.

**Figure 4.**
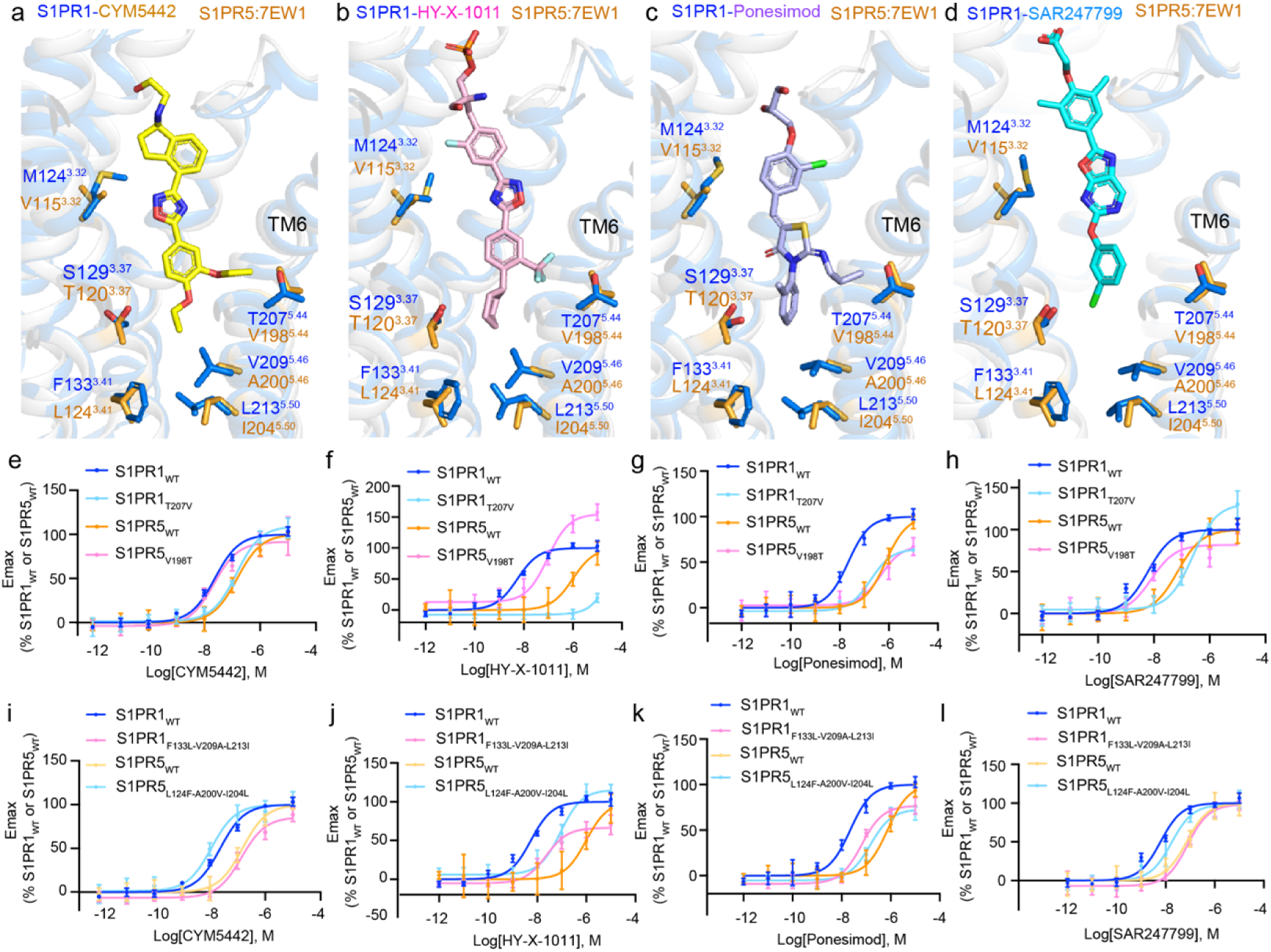
Nonconserved residues influence the selectivity of four agonists for S1PR1 and S1PR5. **A-D.** Nonconserved residues in the ligand binding pocket of S1PR1 and S1PR5. CYM5442-, HY-X-1011-, Ponesimod-, or SAR247799-bound S1PR1 are superimposed with S1PR5(PDB: 7EW1). Residues of S1PR1 (marine) and S1PR5 (orange) are shown as sticks. Molecules are shown as sticks and colored yellow (CYM5442), pink (HY-X-1011), light blue (Ponesimod), and cyan (SAR247799). Orange dashed lines indicate the distances between the agonist and specific residues. Three nonconserved residues located at the bottom of S1PR1, S1PR3, and S1PR5 are enclosed within dashed-line rounded rectangles (orange). **E–L.** BRET-based Gi dissociation assays showing activation of S1PR1 and S1PR5 swapped mutants by the four agonists. Data are presented as mean ± SEM; n=3.

Furthermore, we found that R78^2.37^ in S1PR1, located at the Gi-binding interface and interacting with D350 on the α5 helix of Gi, is not conserved in S1PR5 (A69^2.37^) and S1PR3 (N72^2.37^) (Figure S6A, Supporting Information). Gi dissociation assays using mutant receptors confirmed that R78^2.37^ in S1PR1 and A69^2.37^ in S1PR5 also influence the selectivity of the four agonists for S1PR1 and S1PR5 (Figure S6B–E, Supporting Information).

### 2.4 Roles of non-conserved residues at positions 3.41, 5.46, and 5.50 on agonist subtype selectivity

Most nonconserved residues within the ligand-binding pocket influence receptor subtype selectivity through direct contact with agonists. However, nonconserved residues at positions 3.41, 5.46, and 5.50—two of which do not directly interact with the four agonists in our determined structures—nonetheless play significant roles in determining selectivity for S1PR1, S1PR3, and S1PR5, as revealed by previous analyses. To further understand the structural basis underlying the effects of these nonconserved residues at positions 3.41, 5.46, 5.50, and 5.44 on subtype selectivity, we conducted additional structural analysis.

We observed that residues F/C/L3.41, V/I/A5.46, and L/I/I5.50 are located at the bottom of the orthosteric pockets in S1PR1, S1PR3, and S1PR5, near the intracellular side (Figure 5A–C). Notably, S1PR1 and S1PR5 contain a larger subpocket that accommodates branched functional groups present in the lower regions of agonists. In contrast, S1PR3 features a smaller and shallower subpocket at the corresponding position. Moreover, upon S1PR1 activation, F133^3.41^, V209^5.46^, and L213^5.50^ move inward, leading to stronger interactions between F133^3.41^ and L213^5.50^ and enhancing contacts between TM3 and TM5 (Figure 5D). However, the shorter or smaller side chains of the corresponding residues in S1PR3 (C127^3.41^ and I207^5.50^) and S1PR5 (L124^3.41^ and I204^5.50^) weaken these interactions in those subtypes. Additionally, L/I/I^5.50^ is part of the conserved Class A GPCR activation-related motif P^5.50^I^3.40^F^6.44^, which influence receptor activation.

**Figure 5.**
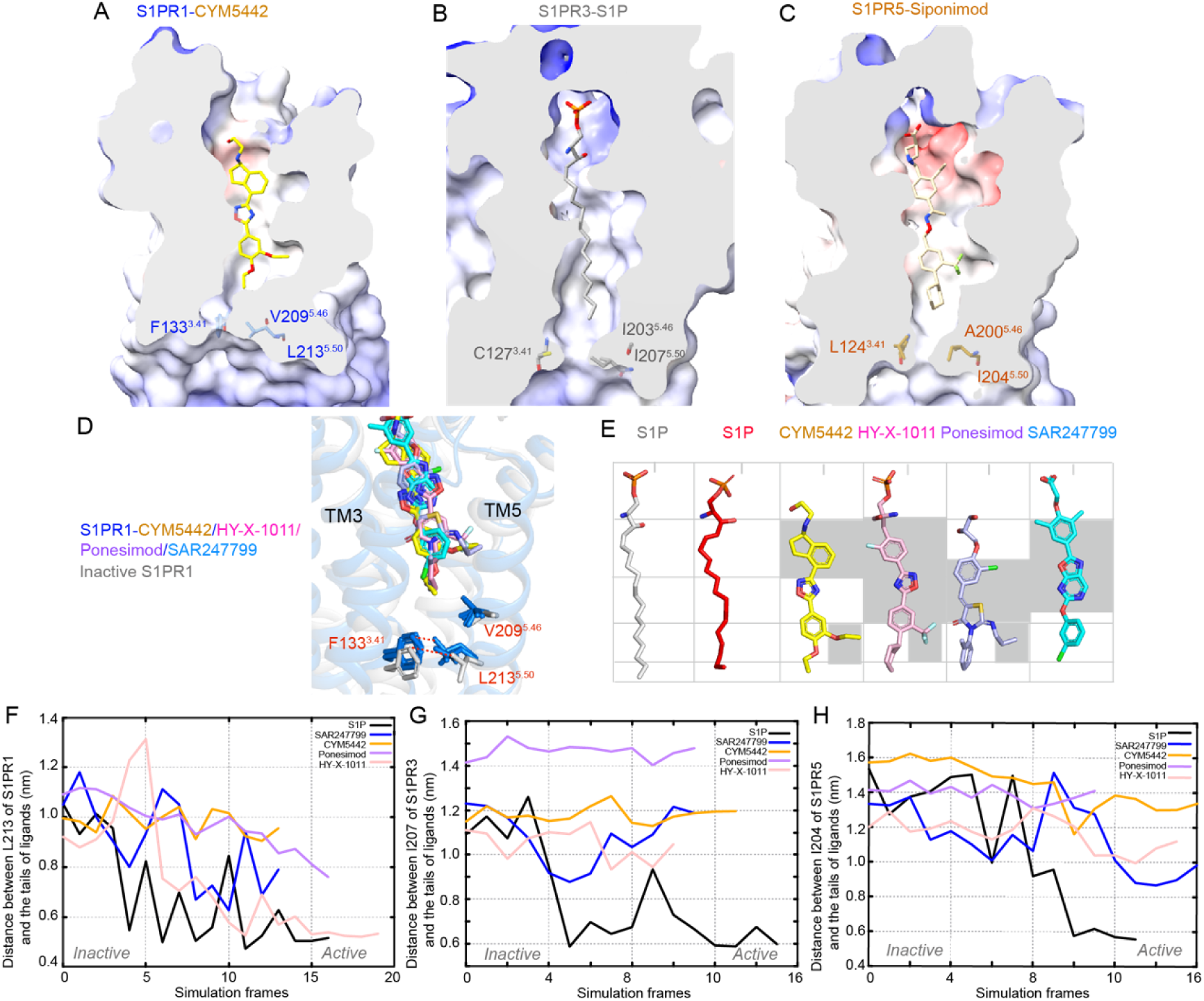
Nonconserved residues at positions 3.41, 5.46, and 5.50 are associated with differences in activation of S1PR1, S1PR3, and S1PR5 induced by five agonists. **A–C.** Three nonconserved residues are located at the bottom of the ligand-binding pockets in S1PR1, S1PR3, and S1PR5. Electrostatic potentials of the pocket and protein surface are displayed: red indicates negative charge, blue indicates positive charge, and white represents hydrophobic regions. The pocket-cutting surface is shown in gray, with all surfaces rendered as semitransparent. Structures include S1PR1 bound to CYM5442 (**A**), S1PR3 bound to S1P (PDB ID: 7EW3) (**B**), and S1PR5 bound to Siponimod (**C**). The bottom portions of these pockets are open toward the intracellular side. Ligands and the three nonconserved residues at the base of each pocket are shown in stick representation. **D.** Superposition of active S1PR1 structures bound to CYM5442, HY-X-1011, Ponesimod, or SAR247799 with the inactive form of S1PR1 (PDB ID: 3V2Y). Residues are shown as sticks, and red lines indicate structural differences between active and inactive conformations. **E.** Structural comparison of the five S1PR1 agonist molecules. Binding modes of the five agonists within the S1PR1 pocket are presented. S1P (gray) corresponds to the structure bound with S1PR3 (PDB ID: 7EW3), while S1P (red) refers to the complex with S1PR1 (PDB ID: 7VIE). The gray background highlights conserved features among four of the molecules. **F–H.** Distances between the tails of CYM5442, HY-X-1011, Ponesimod, or SAR247799 and the Cα atom of residue ^5.50^ in S1PR1, S1PR3, and S1PR5 during receptor activation—from the inactive to active state—calculated using the TAPS method in molecular dynamics simulations.

We hypothesize that S1P, which lacks bulky branched substituents, can easily access the deeper regions of the receptor pocket during activation, thereby facilitating the activation of S1PR3 and S1PR5 and interacting with residues at positions 3.41, 5.46, and 5.50. Compared to S1P, the four synthetic agonists possess larger molecular dimensions (Figure 5E). The branched functional moieties located at the lower portions of CYM5442, HY-X-1011, and Ponesimod appear to hinder these molecules from reaching deeper into the pocket during activation, potentially increasing their selectivity for S1PR1 over S1PR3 and S1PR5. SAR247799, which lacks branched moieties at its lower region, is more likely to penetrate deeply into the S1PR5 pocket. To test this hypothesis, we employed molecular dynamics simulations using our previously established TAPS method to examine agonist binding within the S1PR1/3/5 pockets during the transition from inactive to active receptor states. Distance measurements between the tail ends of the five agonists and residue 5.50 at the bottom of the S1PR1/3/5 pockets supported our hypothesis (Figure 5F–H). Upon receptor activation, S1P penetrates deeply into the pockets of all three subtypes—S1PR1, S1PR3, and S1PR5—whereas CYM5442, HY-X-1011, and Ponesimod reach only relatively shallow positions in the pockets, especially in S1PR3, and S1PR5. SAR247799 inserts more deeply into the S1PR5 pocket compared to CYM5442, HY-X-1011, and Ponesimod, consistent with its dual activity toward S1PR1 and S1PR5.

### 2.5 Structural features and binding modes of agonists critical for enhancing selectivity of S1PR1 over S1PR3 and S1PR5

Although nonconserved residues among S1PR subtypes play crucial roles in ligand selectivity, the designed agonists exhibit greater selectivity for S1PR1 compared to S1P, which demonstrates broad affinity across all five S1PR subtypes. Beyond the above-mentioned structural differences between the four agonists and S1P, their distinct binding modes within the ligand-binding pocket may provide insights into subtype-specific signaling mechanisms mediated by these agonists. Therefore, we further analyzed and compared the binding modes of the four agonists with that of S1P.

Compared to previously reported structures of S1P bound to S1PR1 and S1PR3, the central regions of CYM5442, HY-X-1011, Ponesimod, and SAR247799 adopt a characteristic “curved” conformation, positioning them closer to two nonconserved residues—L^6.55^ and L^7.39^—located at the TM6 and TM7. Structural superposition of S1PR1 with S1PR3 revealed that these four agonists cause steric clashes solely with F263⁶·⁵⁵ or I284⁷·³⁹ or both (Figure 6A–D). Furthermore, compared to S1P-bound S1PR1 and S1PR3, the lower branched moieties of CYM5442, HY-X-1011, and Ponesimod extend deeper into the subpocket formed at the TM5–TM6 interface of S1PR1. This subpocket is composed of F210^5.47^, T207^5.44^, F273^6.52^, and L276^6.55^ in S1PR1, corresponding to F210^5.47^, I201^5.44^, F273^6.52^, and F263^6.55^ in S1PR3 and F210^5.47^, V198^5.44^, F273^6.52^, and FL271^6.55^ in S1PR5 (Figure 6E-L). However, compared to F263⁶·⁵⁵ in the active state of S1PR1, F263⁶·⁵⁵ in S1P-bound S1PR3 adopts a different conformation, significantly reducing the volume of this subpocket and causing steric hindrance with the lower branched moieties of CYM5442, HY-X-1011, and Ponesimod upon structural alignment of S1PR1 with S1PR3 (Figure 6E–H). This likely diminishes the binding affinity of these three agonists toward S1PR3. Similarly, when aligning S1PR1 with S1PR5, compared to Siponimod bound in S1PR5, the lower branched moieties of CYM5442, HY-X-1011, and Ponesimod insert more deeply into the subpocket located at the TM5–TM6 interface of S1PR1. Notably, Siponimod is an approved drug for treating multiple sclerosis (MS) with comparable affinity for both S1PR1 and S1PR5^22,25^. Moreover, the lower branched moieties of these three agonists are close to F210^5.47^, and the nonconserved residue T207⁵·⁴⁴ in S1PR1 corresponds to V198^5.44^ in S1PR5, potentially influencing receptor activation. These structural observations align well with our BRET assay results.

**Figure 6.**
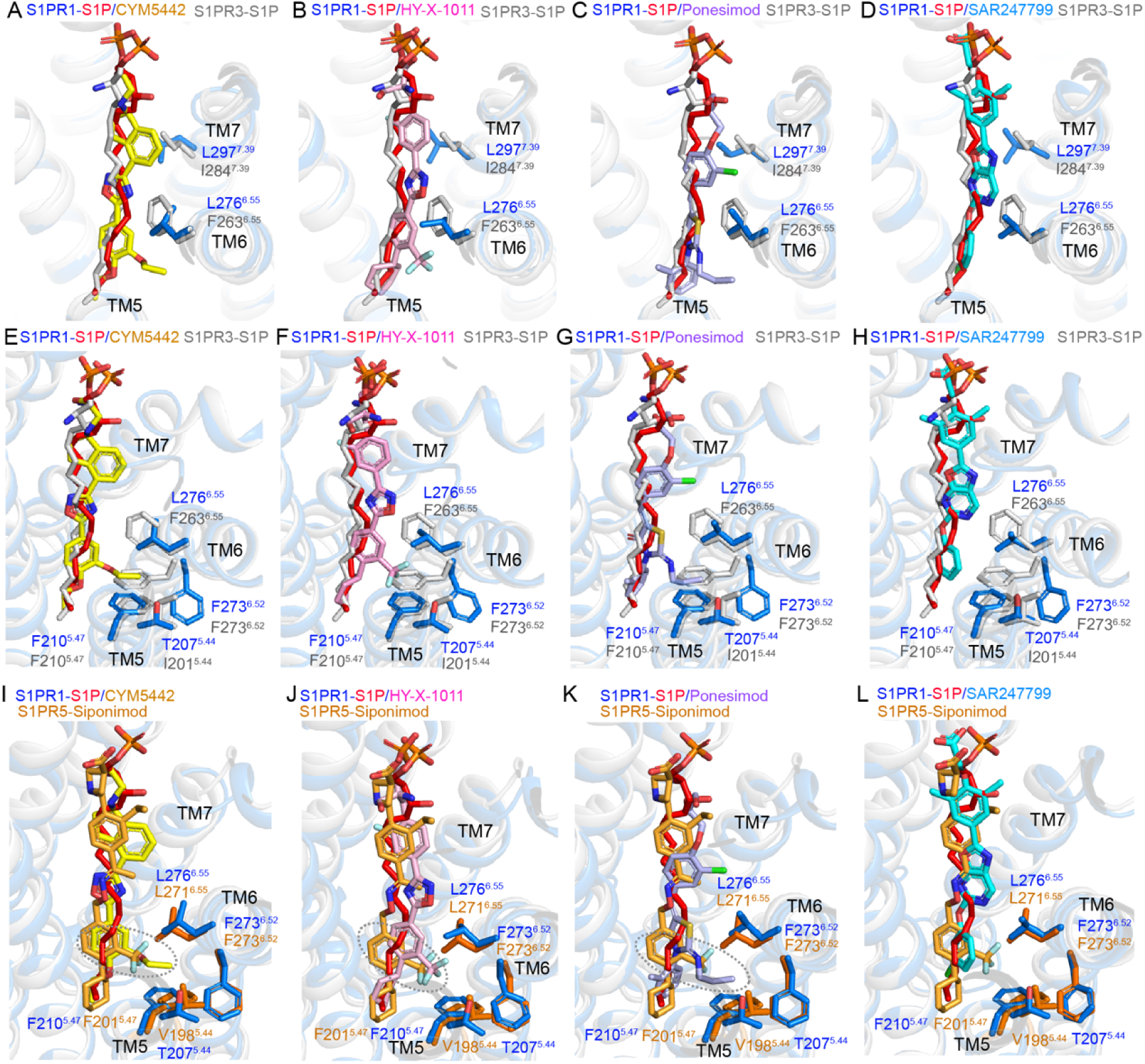
Differences in the binding modes of CYM5442, HY-X-1011, Ponesimod, or SAR247799 bound to S1PR1 compared with S1P bound to S1PR1 and S1PR3, as well as Siponimod bound to S1PR5. **A-H.** Superimposition of CYM5442, HY-X-1011, Ponesimod, or SAR247799 bound to S1PR1 with S1P bound to S1PR1 (PDB: 7VIE) and S1PR3 (PDB: 7EW3). Residues of S1PR1 (marine) and S1PR3 (gray) are shown as sticks. Molecules are displayed as sticks colored as follows: yellow for CYM5442, pink for HY-X-1011, light blue for Ponesimod, cyan for SAR247799, red for S1P (from 7VIE), and gray for Siponimod (from 7EW3). **I-L.** Superimposition of CYM5442, HY-X-1011, Ponesimod, SAR247799, and S1P in S1PR1 (PDB: 7VIE) with Siponimod in S1PR5 (PDB: 7EW1). Four residues of S1PR1 (marine) and S1PR5 (yellow-orange) are shown as sticks. Ligands are represented as sticks and colored as follows: yellow for CYM5442, pink for HY-X-1011, light blue for Ponesimod, cyan for SAR247799, red for S1P (from 7VIE), and yellow-orange for Siponimod (from 7EW1). Dashed gray circles highlight differences in binding poses between Siponimod and CYM5442, HY-X-1011, or Ponesimod.

Based on these findings, we propose that the selectivity of CYM5442, HY-X-1011, Ponesimod, and SAR247799 for S1PR1 over S1PR3 and S1PR5 arises from two key structural features: (1) their “curved” binding orientation toward the TM5–TM7 region, resulting in steric clashes with F263⁶·⁵⁵–I284⁷·³⁹ in S1PR3; and (2) the presence of lower branched moieties that restrict deep insertion into the base of the binding pocket, particularly in S1PR3 and S1PR5.

## 3. Discussion

In this study, we integrated structural analysis, molecular dynamics simulations, and pharmacological assays to elucidate the molecular basis for the selectivity of CYM5442, HY-X-1011, Ponesimod, and SAR247799 toward S1PR1 over S1PR3 and S1PR5. Specifically, selectivity arises from a combination of non-conserved residues in both the ligand-binding pocket and the Gi-protein interaction interface, the distinct curved agonist-binding modes directed toward transmembrane helices 5–7, which lead to steric clashes with two non-conserved residues in S1PR3, and the presence of smaller branched moieties on three agonists that restrict deep insertion into the base of the binding pocket, particularly in S1PR3 and S1PR5. Collectively, these features diminish agonist potency at S1PR3 and S1PR5 and may represent general principles for designing next-generation S1PR1 highly selective agonists. These findings provide a structural framework for the rational development of next-generation S1PR1 agonists with enhanced therapeutic potential.

Additionally, we observed that the EC50 value of HY-X-1011 for S1PR3 is higher than those of CYM5442, Ponesimod, and SAR247799 for the same receptor (Table S3, Supporting Information). The primary distinction between HY-X-1011 and the other three agonists lies in its notably extended structure, allowing for more extensive interactions with residues located in the upper region of the binding pocket. These interactions may contribute to the higher potency of HY-X-1011 toward S1PR3 compared to the other three agonists. Notably, SAR247799 exhibits the weakest potency for S1PR3 among the four agonists, although it is not the shortest molecule. Furthermore, we observed that the tail end of SAR247799 binds at a shallower site compared to CYM5442, HY-X-1011, and Ponesimod within the S1PR1 pocket (Figure 3A-D). The presence of two methyl groups on the benzene ring of SAR247799 restricts its deep insertion into the base of the S1PR1 binding pocket, which may have a more pronounced effect on its binding to S1PR3. Thus, the relatively wide dimension at the upper part of agonists like SAR247799, which restricts their deep insertion into the base of the binding pocket, appears to enhance selectivity for S1PR1 over S1PR3.

Moreover, GPCR signaling via G proteins can be terminated by arrestin binding. Receptor phosphorylation by G protein-coupled receptor kinases (GRKs) promotes arrestin recruitment, which blocks further G protein interaction and initiates receptor internalization. Many S1PR modulators activate both G protein-dependent and arrestin-dependent signaling pathways, which may result in opposing physiological effects. Therefore, developing S1PR modulators with bias toward either G protein or arrestin signaling is crucial for creating more effective and targeted therapeutics targeting S1PR1. SAR247799, a previously reported Gi-biased agonist of S1PR1, exhibits endothelial protective properties and has recently been shown to preserve endothelial barrier integrity in inflammatory bowel disease (IBD) models^37,40^. In our study, we analyzed SAR247799 recognition and activation mechanisms of S1PR1 and initiating downstream Gi signaling. However, the molecular basis for its G protein bias remains unclear. Structural determination of the S1PR1–β-arrestin complex could provide insights into the mechanisms underlying G protein or β-arrestin bias, thereby facilitating the rational design of biased S1PR1-selective agonists with improved therapeutic profiles.

## Methods Section

### ScFv16 Purification

ScFv16 was subcloned into the pFastBac vector and expressed in *Trichoplusia ni* Hi5 insect cells using the Bac-to-Bac system, exactly following the established protocol^33^. Briefly, Hi5 cells were infected by ScFv16 baculovirus (amplified in Sf9 cells) for 96 hours. The medium was collected by centrifugation, pH-balanced to 8.0, and then supplemented with 1 mM NiCl₂ and 5 mM CaCl₂. After incubating for 1h at RT, the supernatant was mixed with Ni-Sepharose resin (GE Healthcare) for another 1 h. The resin was then collected (800×*g*, 10 min, 4°C) and transferred to a gravity column. The column was washed with wash buffer C (20 mM HEPES pH 7.5, 100 mM NaCl, 20 mM imidazole) and eluted with a high-imidazole buffer (wash buffer C containing 250 mM imidazole). HRV-3C protease was added to remove the C-terminal His-tag. Following the tag removal, size-exclusion chromatography was performed on a Superdex 200 Increase 10/300 GL column (GE Healthcare) pre-equilibrated with 20 mM HEPES (pH 7.5) and 100 mM NaCl. The target fractions were concentrated, flash-frozen in liquid nitrogen, and stored at –80°C.

### S1PR1 and G protein Expression and complex Purification

As previously described^33^, the full-length wild-type human S1P1 with the BRIL protein fused to its N-terminus was subcloned into the pFastBac vector. Meanwhile, the wild-type (WT) G_αi1_ and G_β1γ2_ were respectively inserted into the pFastBac vector and pFastBac-Dual vector. The complex was co-expressed in *Spodoptera frugiperda* (Sf9) insect cells with baculoviruses encoding S1P1, WT G_αi1_, and WT G_β1γ2_ at a 1:1:1 multiplicity of infection. Infected cells were harvested after 48 hours, followed by flash-freezing in liquid nitrogen, and stored at –80°C until use. To stabilize the complex, all purification steps were performed in the presence of 10 μM ligands (CYM5422, HY-X-1011, Ponesimod, SAR247799). The cell pellets were thawed and resuspended in lysis buffer (25 mM HEPES pH 7.5, 150 mM NaCl, 5% glycerol, 10 mM MgCl₂, 20 mM KCl, 5 mM CaCl₂, 1 mM MnCl₂, 100 μM PMSF, 2 μg/mL aprotinin, 2 μg/mL pepstatin) and sequentially treated with 25 mU/mL apyrase (1 h, room temperature) and 1% DDM/0.1% CHS (2 hours, 4°C). After centrifugation (39,191 ×g, 30 min, 4°C), the supernatant was incubated with anti-Flag resin (GenScript) in wash buffer A (25 mM HEPES pH 7.5, 150 mM NaCl, 5% glycerol, 5 mM MgCl₂, 5 mM CaCl₂, 0.01% LMNG, 0.001% CHS). Bound proteins were eluted using wash buffer A supplemented with 200 μg/mL Flag peptide. The eluate was further purified by Ni-NTA affinity chromatography. After loading onto Ni-NTA resin (Qiagen), the column was washed with wash buffer B (25 mM HEPES pH 7.5, 150 mM NaCl, 5 mM MgCl₂, 25 mM imidazole, 0.01% LMNG, 0.001% CHS) and eluted with a high-imidazole buffer (wash buffer B containing 250 mM imidazole). The eluted complex was concentrated to less than 2 mL using an Amicon Ultra Centrifugal Filter (MWCO 100 kDa) and incubated with excess scFv16 for 2 hours on ice. Final purification was achieved by size-exclusion chromatography (Superdex 200 Increase 10/300 GL, GE Healthcare) in SEC buffer (25 mM HEPES pH 7.5, 150 mM NaCl, 0.00075% LMNG, 0.00025% GDN, 0.000075% CHS, 100 μM TCEP). Monomeric fractions containing the complex were concentrated to 10-15 mg/mL for cryo-EM grid preparation.

### Molecular cloning of constructs used in BRET assay

The full-length wild-type human S1PR1, S1PR3, S1PR5, and mutants were cloned into a pcDNA3.1 vector with a HA signal peptide, an N-terminal Flag tag. The construct was generated with a standard PCR-based strategy and homologous recombination (1.1×S4 Fidelity PCR Mix, Beijing Genesand Biotech Co., Ltd).

### Cell culture of BRET assay

HEK293T cells (ATCC CRL-11268; mycoplasma free) were maintained, passaged, and transfected in DMEM medium containing 10% FBS, 100 U/ml penicillin, and 100 μg/ml streptomycin (Gibco-ThermoFisher) in a humidified atmosphere at 37°C and 5% CO_2_. After transfection, cells were plated in DMEM containing 2% dialyzed FBS for BRET assay.

### Bioluminescence resonance energy transfer assay (BRET)

BRET assays were used to measure agonists induced S1PR1/3/5 (wild type or mutants) activation coupled Gi signal^44^. HEK293T cells (ATCC CRL-11268; mycoplasma free) were co-transfected in a 1:1:1:1 ratio of receptor: Gα_i1_-Rluc8: G_β_: G_γ_-GFP_2_ with polyethyleneimine. After at least 18 hours, transfected cells were harvested and reseeded in opaque white bottom 96-well assay plates (Beyotime) at a density of 30,000-50,000 cells per well in media (DMEM added 2% dFBS). The next day, the medium was decanted. Cells were incubated in 40 μL 7.5 μM coelenterazine 400a (Goldbio) in drug buffer (1×Hank’s balanced salt solution (HBSS), 20 mM HEPES, pH 7.4, 0.1%BSA) for 2 min, and then treated with 20 μL compounds (CYM5442, HY-X-1011, Ponesimod, and SAR247799) prepared in drug buffer at serial concentration gradient for an additional 5 min. Plates were read in an LB940 Mithras plate reader (Berthold Technologies) with 395-nm and 510-nm emission filters with 1 s per well integration times. BRET ratios were calculated as the ratio of GFP2 emission (510 nm) to Rluc8 emission (395 nm) and analyzed in GraphPad prism 9.0. Ponesimod, CYM5442, and SAR247799 were purchased from MedChemExpress (MCE). HY-X-1011 was synthesized by MCE.

### Cryo-EM sample preparation, data acquisition, and data processing

The purified complex was applied to glow-discharged 300-mesh alloy grids (Quantifoil 300 mesh, Au R1.2/1.3 and M024-Au300-R1.2/1.3) and subsequently vitrified using Vitrobot Mark IV. The images were collected in the counted-Nanoprobe mode on a 300 kV Titan Krios Gi3 electron microscope (Thermo Fisher Scientific) equipped with Gatan K3 Summit detector and GIF Quantum energy filter (slit width 20 eV). All movie stacks with 50 frames were collected using SerialEM software at a nominal magnification of 105,000×, a pixel size of 0.85 Å, and a defocus range of –1.2 μm to –1.8 μm. Each movie stack for S1PR1-Gi-CYM5442, S1PR1-Gi-HY-X-1011, S1PR1-Gi-Ponesimod, and S1PR1-Gi-SAR247799 was recorded for 2.8 s, 2.2 s, 2.3 s, and 2.2 s, respectively, corresponding to a total dose of 53.83 e^-^/Å^2^, 51.73 e^-^/Å^2^, 52.56 e^-^/Å^2^, and 55.63 e^-^/Å^2^, respectively. 3303 movies, 3159 movies, 3141 movies, and 2906 movies were collected for S1PR1-Gi-CYM5442, S1PR1-Gi-HY-X-1011, S1PR1-Gi-Ponesimod, and S1PR1-Gi-SAR247799, respectively.

For S1PR1-Gi-CYM5442, S1PR1-Gi-HY-X-1011, and S1PR1-Gi-Ponesimod data, Data processing was performed using cryoSPARC^45^. Movies frames were aligned using Patch motion. CTF estimation was performed using Patch CTF. Particles were first picked using a blob picker. Particle picking of all micrographs was performed by a template picker. 2,204,113, 3,670,342, and 3,401,413 particles were extracted. Afterwards, particles sets were selected and refined by iterative 2D classification, ab initio reconstruction, and heterogeneous refinement. Finally, 198,838, 1,019,239, and 529,510 particles were selected for S1PR1-Gi-CYM5442, S1PR1-Gi-HY-X-1011, and S1PR1-Gi-Ponesimod data, respectively. After one round of ab initio reconstruction, non-uniform and local refinement, 3.69 Å, 2.79 Å, and 2.79 Å map were refined out for S1PR1-Gi-CYM5442, S1PR1-Gi-HY-X-1011, and S1PR1-Gi-Ponesimod data, respectively.

For S1PR1-Gi-SAR247799 data, movies were imported into RELION 5.0^46^. Beam-induced motion was corrected using MotionCor2^47^, after which the micrographs were imported into CryoSPARC for contrast transfer function (CTF) parameter estimation using PatchCTF. 4,653,961 particles were picked and extracted in a pixel size of 1.7 Å. After 2D classification, 1,364,671 particles were selected for ab-initio reconstruction and heterogeneous refinement. The retained 359,408 particles were re-extracted in RELION in a pixel size of 0.85. These particles underwent additional sieving using CryoSIEVE^48^, resulting in a final selection of 94,216 particles. The final CTF refinement, particle polishing, non-uniform refinement and local refinement result in density map of 3.0 Å resolution.

### Model building and refinement

The initial complex model was built using the structure of S1PR1-Gi-S1P (PDB code: 7VIE) as templates. Models are then fitted into the density map and manually adjusted and fixed in COOT^49^. The restraint files of CYM5442, HY-X-1011, Ponesimod, and SAR247799 were generated by Phenix. elbow package^50^. The complete model was finally refined in Phenix using real-space refinement with secondary structure and geometry restraints^51^ and were checked in COOT. Overfitting of the model was checked by refining the model using one of the two independent maps from gold-standard refinement and calculating FSC against both half maps^52^. The final model was validated using Molprobity^53^ (Table S1). Structural figures were prepared in PyMOL (https://pymol.org/2/), UCSF Chimera^54^, and UCSF ChimeraX^55^.

### Molecular dynamics simulation

AutoDock 4.2^56^ was used to dock the ligands into the binding pocket of the apo inactive/active receptors. In all MD simulations, only receptors and ligands were used, with G protein complex and other protein removed from the structures. The Membrane Builder module in CHARMM-GUI server ^57^ was used to prepare the simulation inputs, a membrane of pre-equilibrated (310 K) POPC lipids based on the OPM database^58^ alignment, TIP3P solvent with 0.15 M Na^+^/Cl^-^ ions and the CHARMM36 force field ^59^. The force field of the ligands was generated by the CGenFF program^60^. All MD simulations were performed using GROMACS-2019.4^61^. The CHARMM36 force-field was used to describe the interactions in the system. Energy minimization was performed for 10000 steps by the steepest descent algorithm and then by the conjugate gradient algorithm. Then a 100 ps NVT simulation was performed at 310 K for solvent equilibration, followed by a 1 ns NPT equilibration to 1 atm using the Berendsen barostat^62^. All MD production simulations were performed with a time-step of 1 fs and a length of 100 ns using Parrinello-Rahman barostat^63^. In all constant temperature simulations, the Bussi (velocity-rescaling) thermostat was used^64^. Long-range electrostatic interactions were treated by the particle-mesh Ewald method^65^. The short-range electrostatic and van der Waals interactions both used a cutoff of 10 Å. All H-bonds were constrained by the LINCS algorithm^66^.

### Path searching

Targeted MD simulations (tMD) were carried out using GROMACS-2019.4 and PLUMED-2.5.3^67^, pulling the S1PR1/3/5-ligands complexes from the inactive states to the active states. The inactive structures were relaxed by a 100-ns MD simulation, and the active structures were energy-minimized, followed by a 100-ps NVT and a 1-ns NPT equilibration. The additional bias potential introduced in tMD takes a simple harmonic form 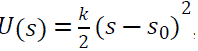, where *k* is the spring force constant, *s*_0_ represents the target structure. *k* was set to 1000 kJ/(mol Å^2^), biasing on heavy atoms of the non-loop parts of receptors and the ligands, while structure alignments were performed using C-alpha atoms of the non-loop parts of receptors.

To find the minimum free energy paths (MFEPs), which are more physically natural than the initial paths, the TAPS method^68^ was used to do path optimization. The TAPS simulation was performed using an in-house python script incorporating GROMACS, PLUMED and Concorde^69^. The simulations were performed in the NVT ensemble at 310 K, using the velocity-rescale thermostat^64^. The initial path was obtained by selecting conformations with a gap of 1.0 Å from the tMD trajectory. The RMSD between the conformations were computed using all heavy atoms of the non-loop parts of receptors and ligands, while structure alignments were performed using C-alpha atoms of the non-loop parts of receptors. In each TAPS iteration, 1000 ps sampling was performed in total. Gaussians of height 0.25 kJ/mol and width 0.5 were deposited every 0.01 ps, with frames recorded at the same frequency. After the optimization, convergence was evaluated by Multidimensional Scaling (MDS) method^70^ and PCV-z analysis^71^.

### Free energy calculation

To evaluate the probability of receptor activation, umbrella sampling ^72^ was used to do the free energy calculation. Umbrella sampling was performed using GROMACS-2019.4 and PLUMED-2.5.3. The free energy profiles of the MFEPs were calculated along PCV-s, which represents the progress along the MFEP. The sampling in each window was restrained within 1.5 Å of MFEP through a harmonic wall potential with a force constant of 20,000 kJ/(molÅ^4^) at PCV-z=2.25Å^2^. The RMSD among the conformations, which was computed by all heavy atoms of the non-loop parts of receptors and ligands, was chosen as the CV. Structure alignment was performed using C-alpha atoms of the non-loop parts of receptors. The window size was chosen as 0.5 Å along the MFEP. For each window, a force constant of 200 kJ/mol was employed. Each window was simulated for at least 10 ns. A complete free energy profile was evaluated with the weighted histogram analysis method (WHAM)^73^. When the iteration process of WHAM reached convergence, the mean value and standard statistical error (error bar) of the free energy were calculated in each window from the last 10 WHAM iterations.

## Data availability statement

Cryo-EM maps and atomic models have been deposited in the Electron Microscopy Data Bank under accession codes: EMD-65218, EMD-65219, EMD-65220, EMD-65221 and in the Protein Data Bank under accession codes: 9VNY, 9VNZ, 9VO0, 9VO1.

## Acknowledgments

We thank the Kobilka Cryo-EM Center at the Chinese University of Hong Kong, Shenzhen for supporting EM data collection. This work is funded by China Postdoctoral Science Foundation (2023M730692).

## Author Contributions

R.B.R. and L.Y.Y. designed and supervised the entire project. L.Y.Y. and H.Z.J. prepared the grids, collected, and processed the cryo-EM data, and built the protein atomic model. B.P. conducted all BRET assays and processed the corresponding data. R.J.T. performed the molecular dynamics simulations and related calculation. B.G. carried out protein expression and purification. Z.Y.Q. conducted molecular cloning of the swapped mutants. J.X.W. provided partial compounds. Z.Y.Z. supervised the molecular dynamics simulation work. H.L.H. assisted with structure validation. L.Y.Y. and R.B.R. wrote and revised the manuscript. B.P., H.Z.J., R.J.T., and B.G. contributed to manuscript preparation.

## Supporting Information

**Figure S1.**
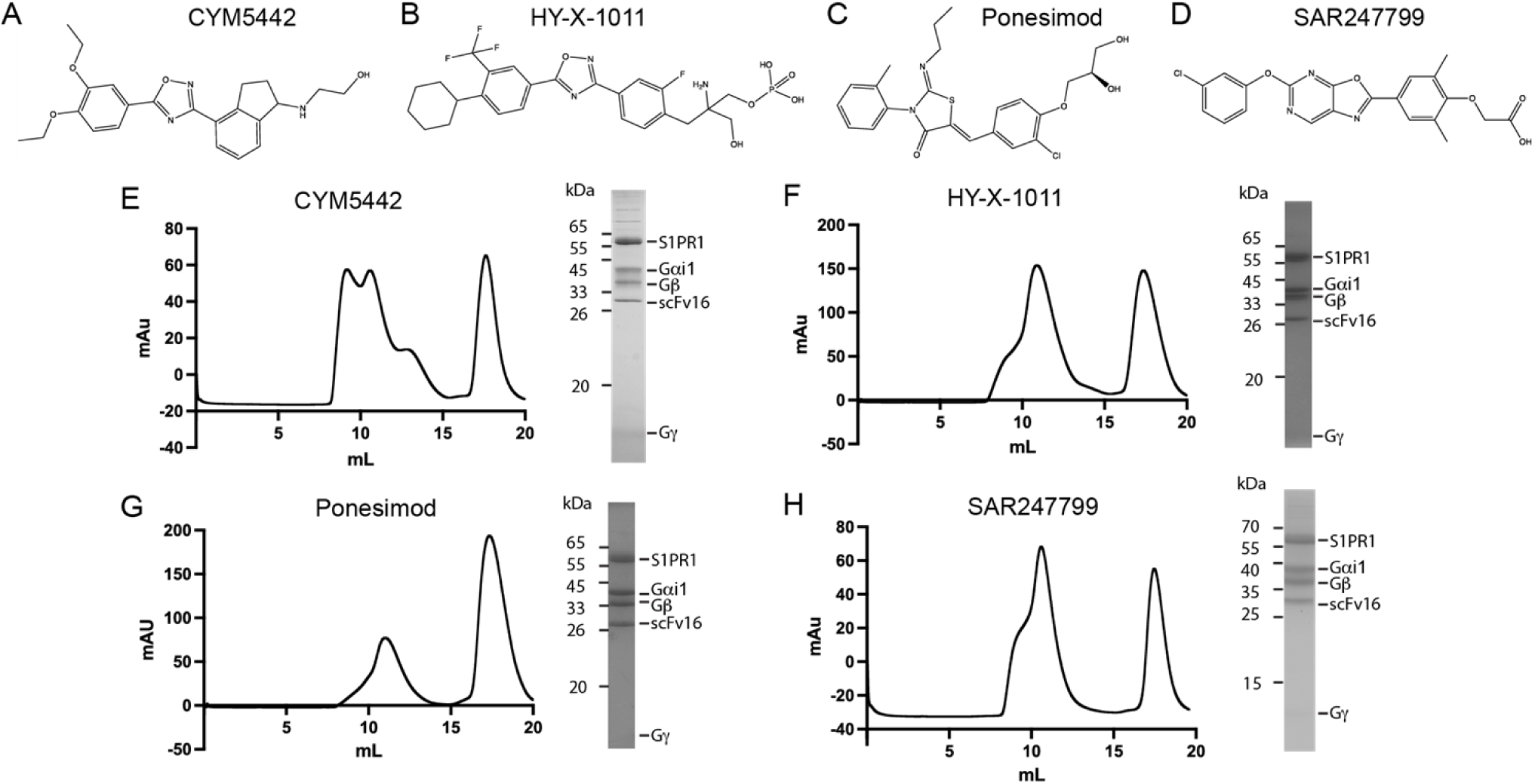
Agonist structure and purification of the S1PR1-Gi complex bound with CYM5442, HY-X-1011, Ponesimod, and SAR247799. **A-D.** Structural formulas of four agonist molecules: CYM5442, HY-X-1011, Ponesimod, and SAR247799. **E-H.** Gel filtration chromatography elution profile of the four complexes and SDS-PAGE of the eluted complex.

**Figure S2.**
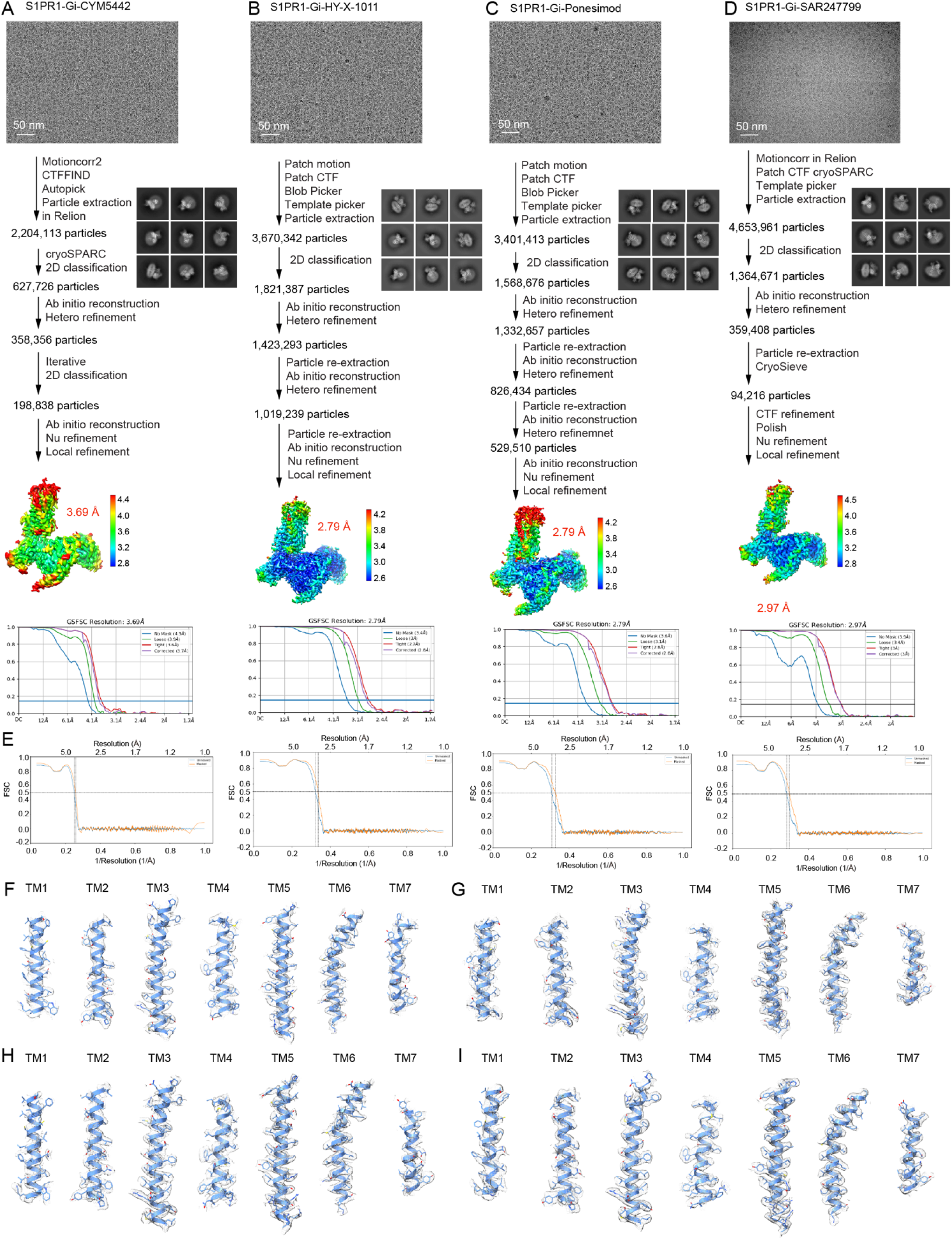
Structural determination of the S1PR1-Gi complex bound with CYM5442, HY-X-1011, Ponesimod, and SAR247799. **A-D.** Flowchart for processing cryo-EM data of S1PR1-Gi bound with CYM5442, HY-X-1011, Ponesimod, or SAR247799. Representative micrographs, 2D classes, local resolution of the final map estimated via cryoSPARC, and Fourier shell correlation (FSC) curves of the final refined cryo-EM map are displayed. **E.** Fourier shell correlation (FSC) between the map and model of the four complexes. FSC curve of the final refined model against the full map. **F-I.** Density maps of the transmembrane helix of S1PR1 in the four complexes are shown as meshes.

**Figure S3.**
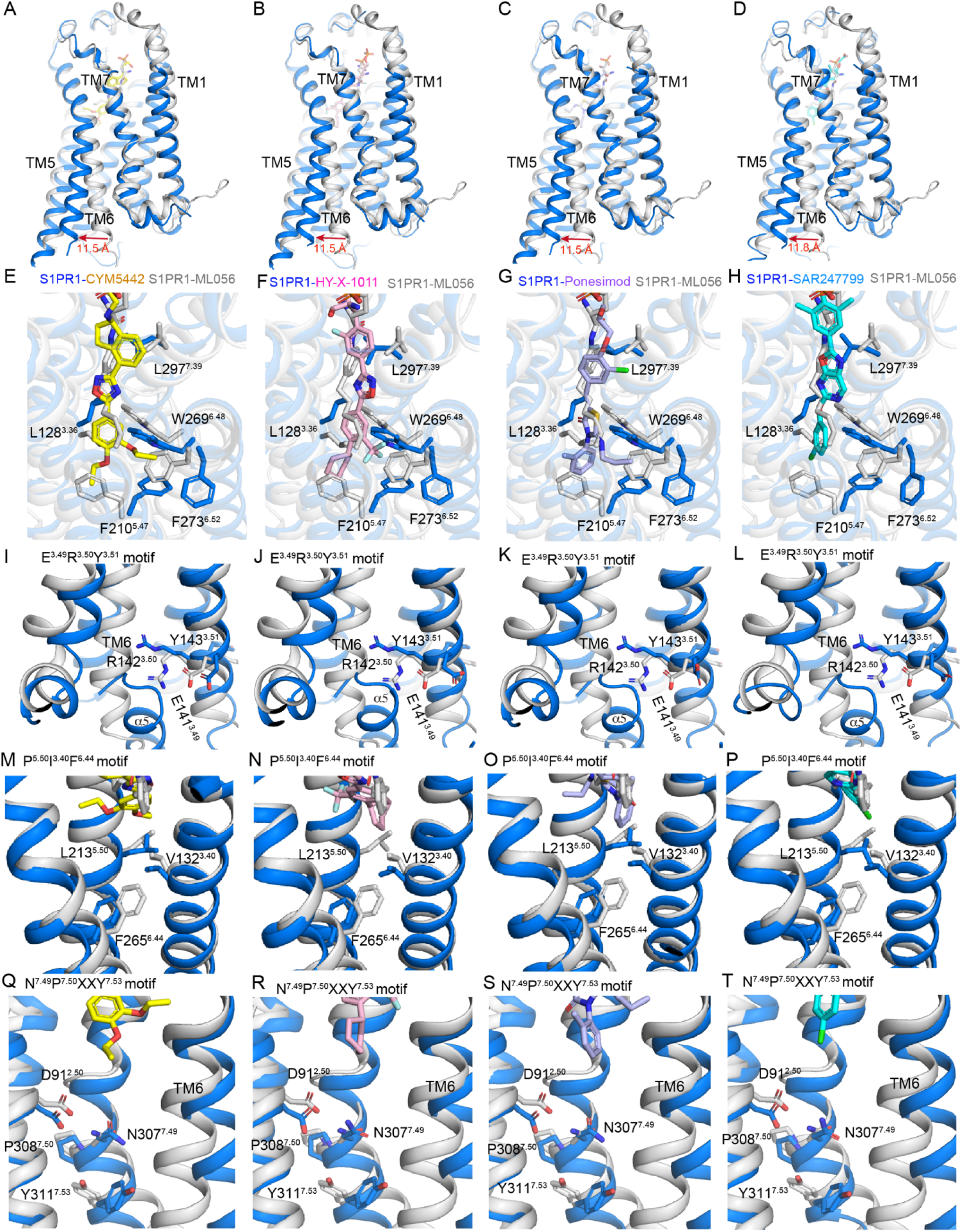
Comparison of the active conformation of S1PR1 activated by CYM5442, HY-X-1011, Ponesimod, and SAR247799 with the inactive conformation of S1PR1. **A-D.** The transmembrane (TM) helix shifts between the active form of S1PR1 (marine-colored) bound with CYM5442, HY-X-1011, Ponesimod, or SAR247799 and the inactive form of the S1PR1–ML056 complex (gray; PDB ID: 3V2Y). Red arrows indicate notable shifts in TM6, with corresponding distance measurements. **E-H.** Conformational changes in key residues at the bottom of the pockets between the active form S1PR1 bound with CYM5442, HY-X-1011, Ponesimod, or SAR247799 and the inactive form S1PR1. Residues and molecules are shown as sticks and colored yellow (CYM5442), pink (HY-X-1011), light blue (Ponesimod), cyan (SAR247799), and gray (ML056). **I-L, M-P, Q-T.** Conformational changes in the E^3.49^R^3.50^Y^3.51^ motif, P^5.50^I^3.40^F^6.44^ motif, and N^7.49^P^7.50^XXY^7.53^ motif between the active form of S1PR1 and CYM5442, HY-X-1011, Ponesimod, or SAR247799 with the inactive form of S1PR1.

**Figure S4.**
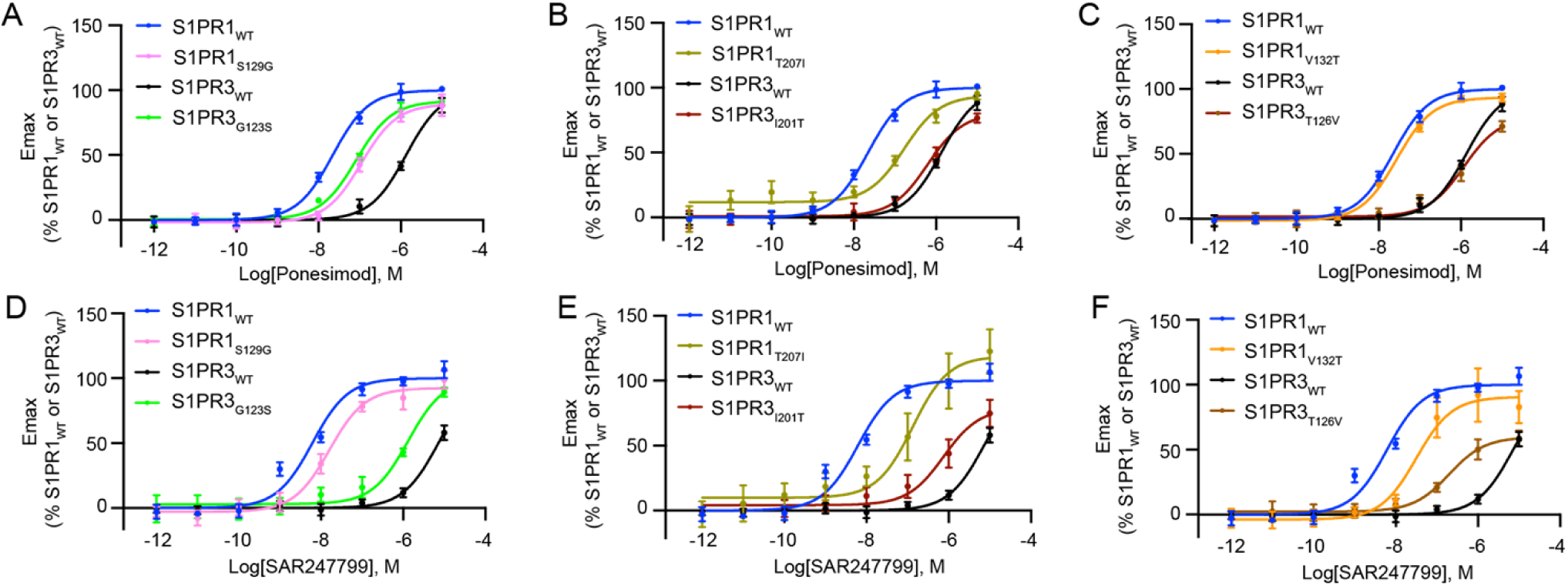
Nonconserved residues influence agonists selectivity for S1PR1 and S1PR3. **A-F.** BRET-based Gi dissociation assays showing activation responses of S1PR1 and S1PR3 swapped mutants (S1PR1_S129G_, S1PR3_G123S_, S1PR1_T207I_, and S1PR3_I201T_) induced by Ponesimod or SAR247799. Data are presented as mean ± SEM; n=3.

**Figure S5.**
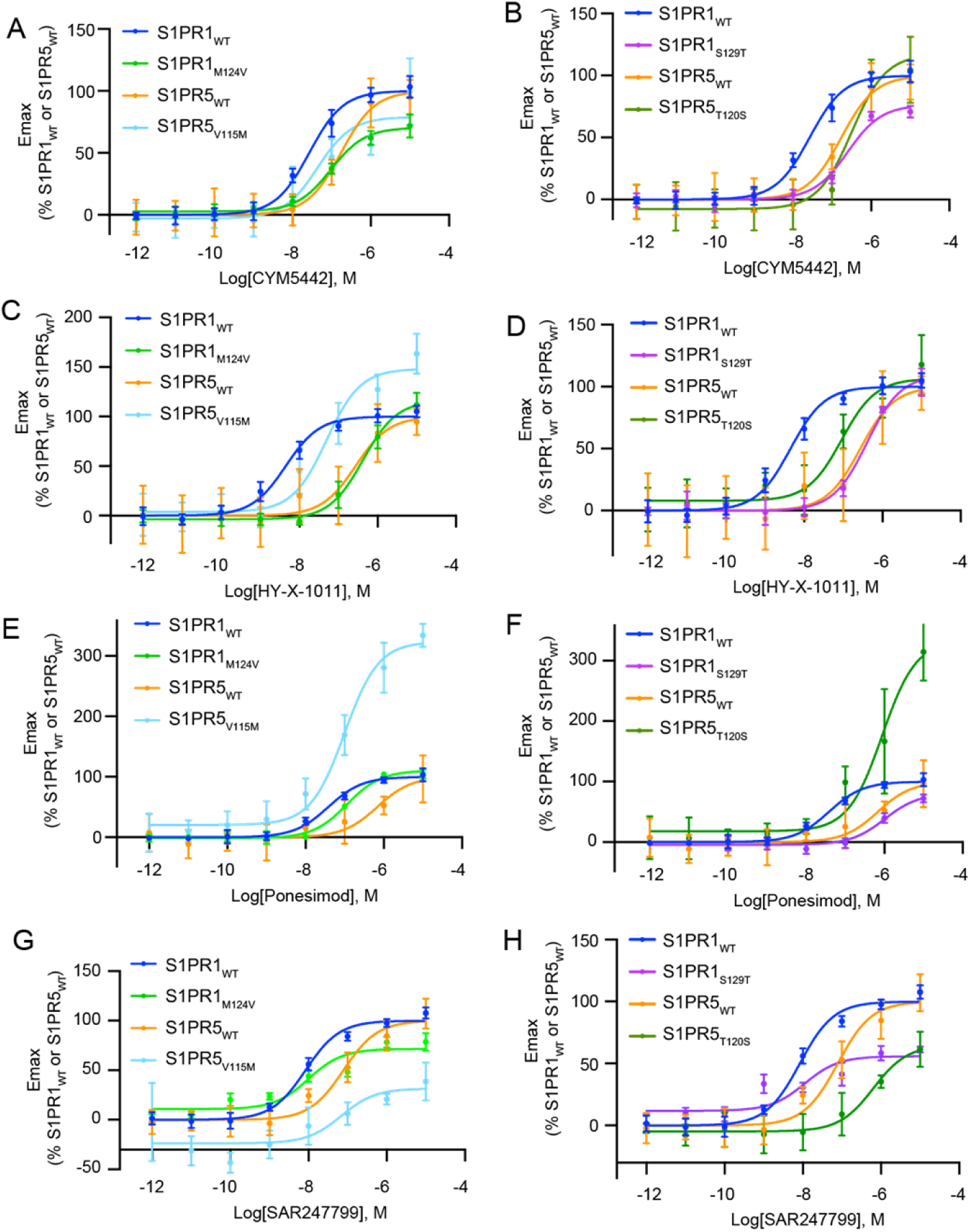
Nonconserved residues influcence four agonists selectivity for S1PR1 and S1PR5. **A-B. C-D. E-F. G-H.** BRET G_i_ dissociation assay of S1PR1 and S1PR5 swapped mutants (S1PR1_M124V_, S1PR5_V115M_, S1PR1_S129T_, and S1PR5_T120S_) activation responses induced by CYM5442, HY-X-1011, Ponesimod, or SAR247799. The data are presented as the means ± SEMs; n=3.

**Figure 6.**
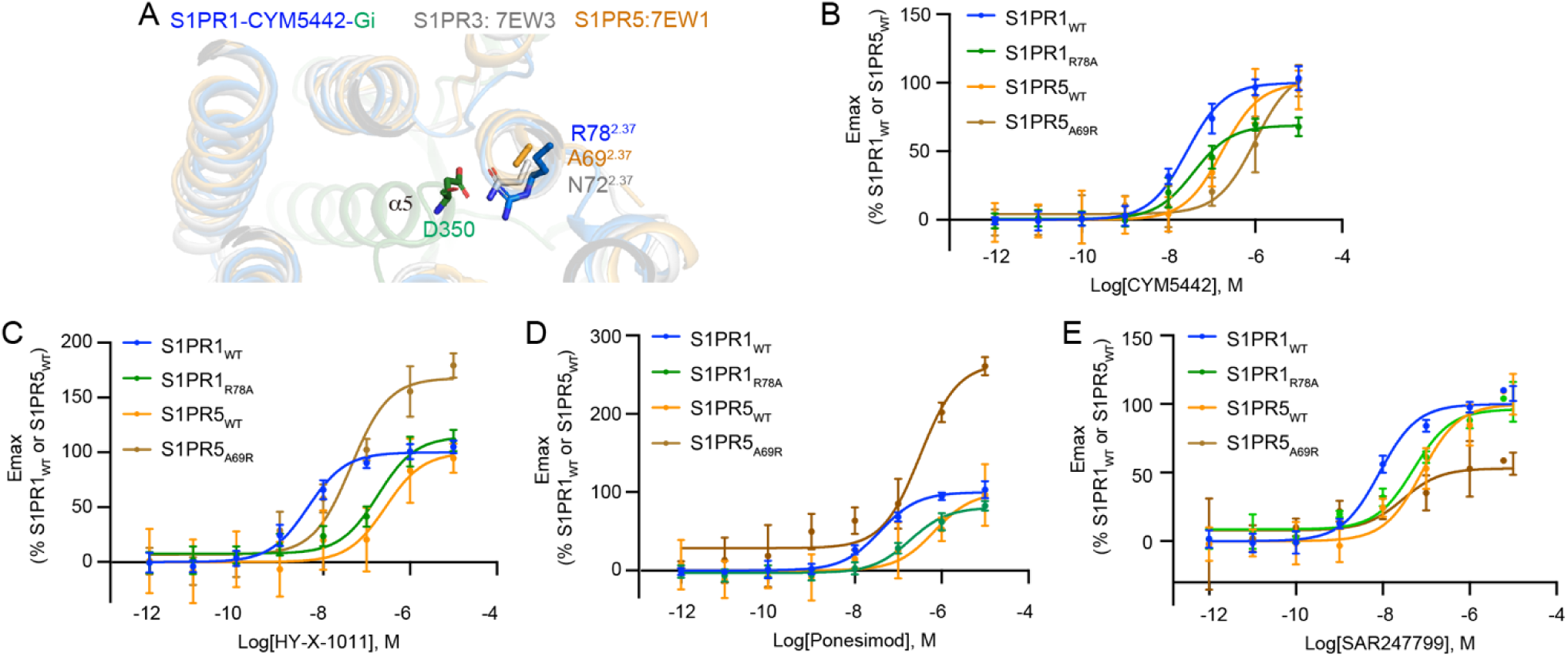
Nonconserved residues at the Gi binding interface of S1PR1 and S1PR5 that influence activation by CYM5442, HY-X-1011, Ponesimod, and SAR247799. **A.** Superimposition of the Gi binding surface of S1PR1-Gi bound with CYM5442, S1PR3 (PDB: 7EW3), and S1PR5 (PDB: 7EW1). Nonconserved residues of S1PR1 (marine), S1PR3 (gray), and S1PR5 (yellow-orange) and D350 of Gi are shown as sticks. Gi is colored in green. **B-E.** BRET Gi dissociation assay showing activation responses of S1PR1 and S1PR5 swapped mutants (S1PR1_R78A_ and S1PR5_A69R_) induced by CYM5442, Ponesimod, or SAR247799. The data are presented as the means ± SEMs; n=3.

**Supplementary table 1.** Cryo-EM data collection, refinement and validation statistics.

|  | S1PR1-CYM5442<br>(EMDB-65218)<br>(PDB 9VNY) | S1PR1-HY-X-1011<br>(EMDB-65219)<br>(PDB 9VNZ ) | S1PR1-Ponesimod<br>(EMDB-65220)<br>(PDB 9VO0) | S1PR1-SAR247799<br>(EMDB-65221)<br>(PDB 9VO1) |
| --- | --- | --- | --- | --- |
| **Data collection and processing** |  |  |  |  |
| Magnification | 105,000 | 105,000 | 105,000 | 105,000 |
| Voltage (kV) | 300 | 300 | 300 | 300 |
| Electron exposure (e-/Å<sup>2</sup>) | 53.83 | 51.73 | 52.56 | 55.63 |
| Defocus range (μm) | -1.2- -1.8 | -1.2- -1.8 | -1.2- -1.8 | -1.2- -1.8 |
| Pixel size (Å) | 0.85 | 0.85 | 0.85 | 0.85 |
| Symmetry imposed | C1 | C1 | C1 | C1 |
| Initial particle images (no.) | 2,204,113 | 3,670,342 | 3,401,413 | 4,653,961 |
| Final particle images (no.) | 198,838 | 1,019,239 | 529,510 | 94,216 |
| Map resolution (Å) | 3.69 | 2.79 | 2.79 | 3.0 |
| FSC threshold | 0.143 | 0.143 | 0.143 | 0.143 |
| Map resolution range (Å) | 2.8-4.4 | 2.6-4.2 | 2.6-4.2 | 2.8-4.5 |
| **Refinement** |  |  |  |  |
| Initial model used (PDB code) | 7VIE | 7VIE | 7VIE | 7VIE |
| Model resolution (Å) | 3.7/3.9 | 2.8/3.0 | 2.8/3.0 | 2.8/3.0 |
| FSC threshold | 0.143/0.5 | 0.143/0.5 | 0.143/0.5 | 0.143/0.5 |
| Model composition |  |  |  |  |
| Non-hydrogen atoms | 8920 | 8931 | 8926 | 8931 |
| Protein residues | 1138 | 1135 | 1136 | 1134 |
| Ligands | 1 | 1 | 1 | 1 |
| *B* factors (Å<sup>2</sup>) |  |  |  |  |
| Protein | 98.51 | 78.34 | 111.84 | 135.91 |
| Ligand | 124.24 | 150.33 | 142.5 | 181.76 |
| R.m.s. deviations |  |  |  |  |
| Bond lengths (Å) | 0.004 | 0.005 | 0.004 | 0.003 |
| Bond angles (°) | 0.840 | 0.667 | 0.677 | 0.591 |
| Validation |  |  |  |  |
| MolProbity score | 1.98 | 1.47 | 1.82 | 1.88 |
| Clashscore | 13.32 | 6.35 | 9.16 | 7.23 |
| Poor rotamers (%) | 0.21 | 0.51 | 0.41 | 1.33 |
| Ramachandran plot |  |  |  |  |
| Favored (%) | 95.08 | 97.40 | 95.25 | 94.34 |
| Allowed (%) | 4.56 | 2.60 | 4.48 | 5.66 |
| Disallowed (%) | 0.36 | 0.00 | 0.27 | 0 |

**Supplementary table 2.** The EC50 and Emax of the S1PR1 mutants activated by four agonist molecules: CYM5442, HY-X-1011, Ponesimod, and SAR247799 in BRET Gi dissociation assay.

| CYM5442 |  |  | HY-X-1011 |  |  | Ponesimod |  |  | SAR247799 |  |  |
| --- | --- | --- | --- | --- | --- | --- | --- | --- | --- | --- | --- |
| Mutants | EC50(nM) | Emax(%) | Mutants | EC50(nM) | Emax(%) | Mutants | EC50(nM) | Emax(%) | Mutants | EC50(nM) | Emax(%) |
| S1PR1<sup>a</sup> | 26.70±0.06 | 100.00 | S1PR1<sup>a</sup> | 7.33±0.07 | 100.00 | S1PR1<sup>a</sup> | 38.09±0.06 | 100.00 | S1PR1<sup>a</sup> | 5.20±0.03 | 100.00 |
| R120A | 227.70±0.07 | 72.65 | N101A | 100.20±0.14 | 42.13 | F125A | 641.60±0.10 | 70.83 | N101A | 81.99±0.50 | 14.51 |
| E121A | 348.00±0.07 | 104.20 | M124A | - | - | L128A | 310.90±0.15 | 69.69 | T109A | 689.50±0.21 | 72.75 |
| T207A | 352.90±0.08 | 63.44 | F125A | 16830.00±0.79 | - | L195A | 246.50±0.07 | 99.47 | E121A | 72.83±0.71 | 15.02 |
| E294A | 322.30±0.08 | 80.90 | L128A | 824.70±0.09 | 104.80 | T207A | 187.40±0.04 | 139.20 | F125A | 1172.00±0.35 | 18.21 |
| S1PR1<sup>b</sup> | 14.78±0.10 | 100.00 | S129A | 371.10±0.15 | 57.91 | W269A | 1281.00±0.05 | 105.00 | L128A | 8.42±0.32 | 22.04 |
| N101A | 1170.00±0.19 | 76.99 | F210A | 69.59±0.18 | 50.73 | S1PR1<sup>b</sup> | 30.27±0.12 | 100.00 | L195A | - | - |
| F125A | 3882.00±0.55 | 66.99 | L276A | 6956.00±0.80 | 59.28 | V132A | 72.34±0.12 | 110.20 | F210A | 89.68±0.09 | 96.20 |
| L128A | 1675.00±0.12 | 113.30 | E294A | 1193.00±0.11 | 116.80 | V194A | - | - | L272A | - | - |
| V132A | 30.52±0.13 | 111.50 | L297A | 1181.00±0.23 | 48.54 | C206A | 225.20±0.12 | 130.20 | L276A | 1164.00±0.21 | 36.39 |
| V194A | 39.17±0.13 | 92.49 | S1PR1<sup>b</sup> | 4.45±0.07 | 100.00 | F210A | 1466.00±0.85 | 25.25 | S1PR1<sup>b</sup> | 8.40±0.06 | 100.00 |
| F210A | 16250.00±0.92 | - | Y29A | - | - | F265A | 1384.00±0.08 | - | R120A | 160.20±0.24 | 20.83 |
| W269A | 1121.00±0.13 | 100.40 | E121A | - | - | L272A | 25.09±0.15 | 92.55 | S129A | 1924.00±1.17 | 9.83 |
| L272A | 4456.00±0.35 | 95.67 | L272A | - | - | L276A | 735.10±0.17 | 85.23 | L297A | 560.80±0.07 | 150.10 |
| L276A | 1425.00±0.17 | - |  |  |  | E294A | 2154.00±0.12 | - | W269A | 382.30±0.24 | 45.98 |
| E294A | 297.70±0.24 | 59.46 |  |  |  | L297A | 1742.00±0.19 | 98.83 |  |  |  |
| L297A | 1387.00±0.38 | 51.50 |  |  |  | S1PR1<sup>c</sup> | 38.09±0.06 | 100.00 |  |  |  |
| S1PR1<sup>c</sup> | 26.70±0.06 | 100.00 |  |  |  | N101A | 87.95±0.12 | 61.15 |  |  |  |
| M124A | - | 12.93 |  |  |  | E121A | 369.60±0.28 | 30.20 |  |  |  |
| S129A | 1633.00±0.27 | 41.63 |  |  |  | S129A | 68.83±0.33 | 35.08 |  |  |  |

a, b, c, d represent the normalized data of the same group respectively

**Supplementary table 3.**
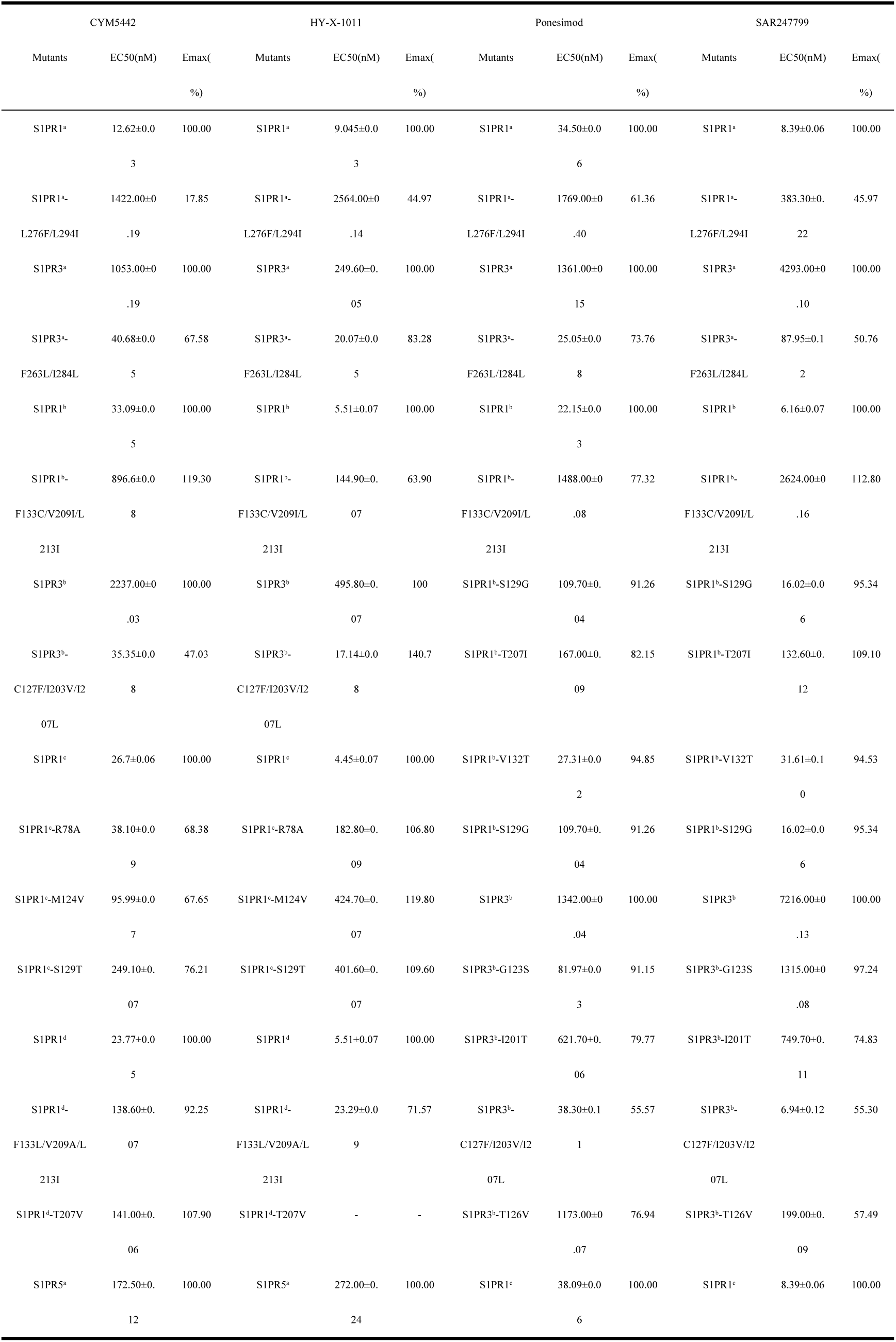
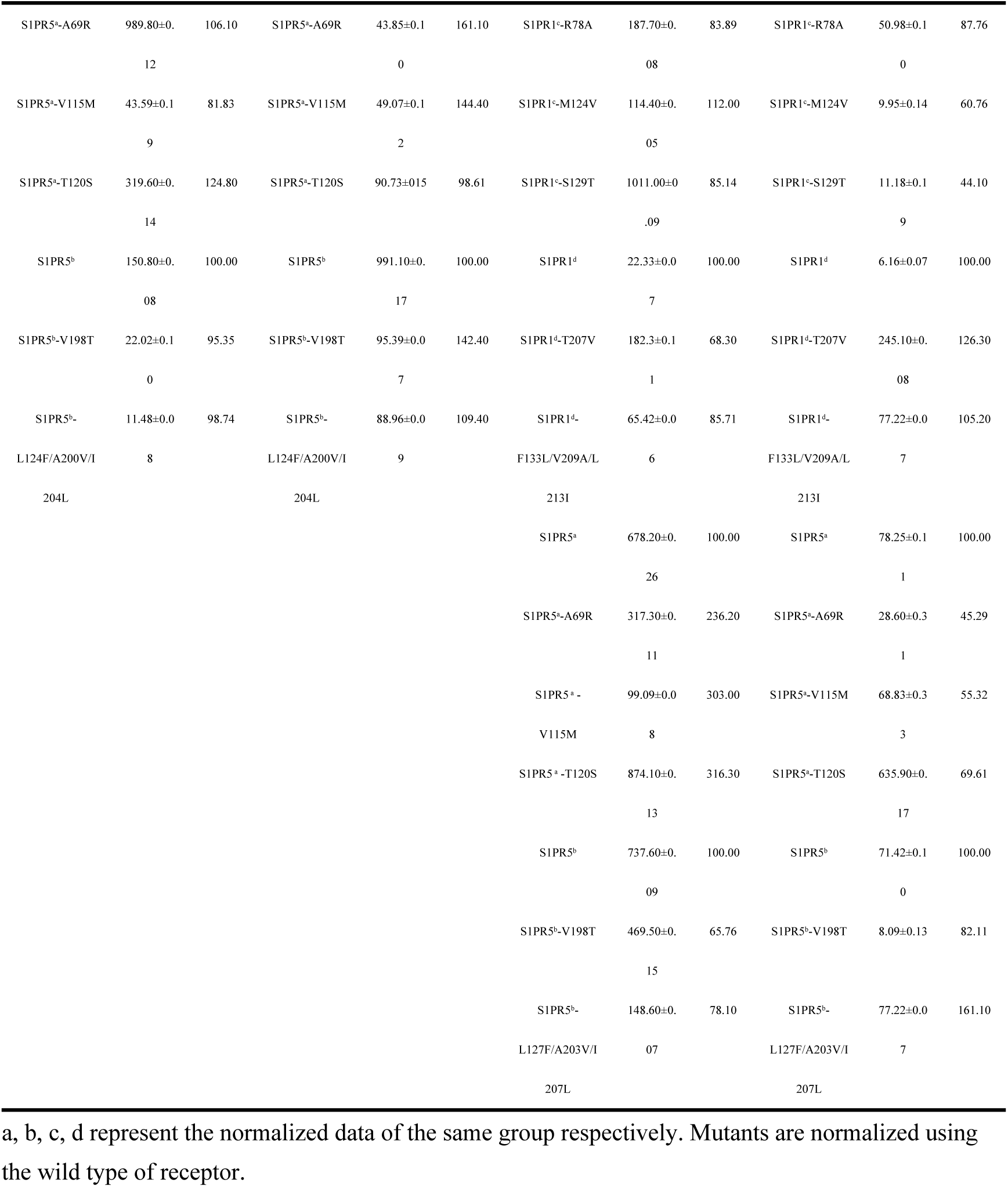
The EC50 and Emax in BRET Gi dissociation assay of swapped mutants of S1PR1, S1PR3, and S1PR5 activated by four agonist molecules: CYM5442, HY-X-1011, Ponesimod, and SAR247799.

